# Metastable Intermediates Identified in Epithelial to Mesenchymal Transition are Regulated by G-Quadruplex DNA Structures

**DOI:** 10.1101/2023.08.21.554220

**Authors:** Jessica J. King, Cameron W. Evans, Ulrich D. Kadolsky, Marck Norret, Munir Iqbal, Clémentine Mercé, Sugandha Bhatia, Philip A. Gregory, Erik W. Thompson, Alka Saxena, K. Swaminathan Iyer, Nicole M. Smith

## Abstract

Cancer is a heterogenous disease, with multiple cellular subpopulations present within a single tumour mass that differ genetically and morphologically, and thus respond differently to chemotherapeutics. Epithelial-to-Mesenchymal transition (EMT) has been shown to play a role in tumour heterogeneity. Single-cell sequencing is critical to identify cell-type-specific transcriptomic differences with multiplexing methods increasing experimental scope with reduced cost. Cell hashing with barcoded antibodies is commonly used to multiplex samples but is limited to samples expressing target antigens. Antigen-independent methods of barcoding cells, such as barcoded lipid-anchors, have gained traction but present substantial populations that cannot be unambiguously demultiplexed. Herein we report a multiplexed single-cell transfection-enabled cell hashing sequencing (scTECH-seq) platform, which uses antigen-independent endocytic uptake to barcode cells, resulting in efficient, uniform barcoding with high cell recovery. We apply this methodology to identify distinct metastable cell states in human mammary cells undergoing EMT and show that stabilisation of G-quadruplex DNA has the potential to inhibit EMT.

---

Epithelial to mesenchymal transition (EMT) is a highly dynamic process in which epithelial cells acquire mesenchymal characteristics via a complex network of cellular changes^1–4^. In the context of cancer, cells undergoing EMT lose cell-to-cell adhesions^5,6^ and develop increased invasiveness, resistance to apoptosis, and migratory properties that facilitate entry into circulation and invasion of foreign tissue^7,8^. The complex programme of gene expression changes during EMT means that individual cells within a population exist along a spectrum and can display epithelial, mesenchymal, as well as intermediate transcriptomic profiles with both epithelial and mesenchymal characteristics^9,10^. These intermediate (or ‘hybrid’) cells can represent minor fractions of the total population but display high invasive potential and unregulated proliferation, characteristic of cancer stem cells^10–12^. EMT therefore presents a major obstacle to effective cancer therapy because it represents a source of circulating tumour cells that contribute to metastasis and the evolution of drug resistance^4,13,14^. The ability to resolve gene expression profiles of multiple hybrid cells in EMT are lost using bulk RNA sequencing methods. Therefore, to fully capture the heterogeneity of EMT samples, high-resolution technologies like single-cell sequencing are required.

Multiplexing samples have extended the scope of single-cell sequencing studies by substantially reducing the cost per sample, allowing sequencing of thousands of samples in parallel. Multiplexed single-cell sequencing methods such as Cell hashing and 10x Genomics’ Cell Plex, utilise DNA barcodes to tag samples prior to pooling. Cell hashing methods typically use barcoded antibodies that target cell membrane proteins, however these methods rely on the ubiquitous expression of the target antigens^15^. The use of antigen-independent universal insertion methods of barcoding the cells, such as lipid-based delivery, have also gained traction however, can give rise to substantial populations that cannot be unambiguously demultiplexed^16,17^. Furthermore, commercially available multiplex methods, including Cell Plex are only available for 3’ Gene expression profiling thus limiting it’s application in a more diverse context. Multiplexed single-cell investigation of samples containing minor cell subpopulations require a near-quantitative cell hashing efficiencyfor accurate deconvolution. Furthermore, the hashing method needs to label cells without bias to avoid significant data exclusion.

We have developed an unbiased multiplex method for single-cell profiling on the 10x Genomics platform, called single cell transfection enabled cell-hashing sequencing (scTECH-seq). Our method uses whitelisted feature barcode sequences for 5’ Gene expression kit provided by 10x Genomics, to create short barcoded oligonucleotides (SBOs). We show that our SBOs label every cell within a given sample and therefore barcode sequences can be used to deconvolute sequenced data and map cells back to their originating samples.

Using scTECH-seq, we identify multiple cellular states expanding the entire spectrum of EMT in human mammary epithelial cells. Furthermore, we find that DNA G-quadruplex structures regulate the expression of EMT driving genes and pathways, and can be targeted to inhibit overall EMT.

## Results

### scTECH-seq has high transfection efficiency

scTECH-seq uses SBOs from the whitelisted feature barcode sequences for the 10x Genomics 5’ Gene expression (GEX) kit. The full sequence structure of SBO is shown in Supplementary Fig. S1. In scTECH-seq, each sample is incubated with a unique SBO, before samples are pooled together, in the required ratios to make a single cell suspension of individually barcoded cells from multiple samples. These barcoded cells are loaded into a single channel of a Chromium chip (Fig. 1a). Single cell libraries are prepared using the standard vendor recommended 5’ GEX protocol, and SBO library is generated using the feature barcode library protocol.

**Fig. 1.**
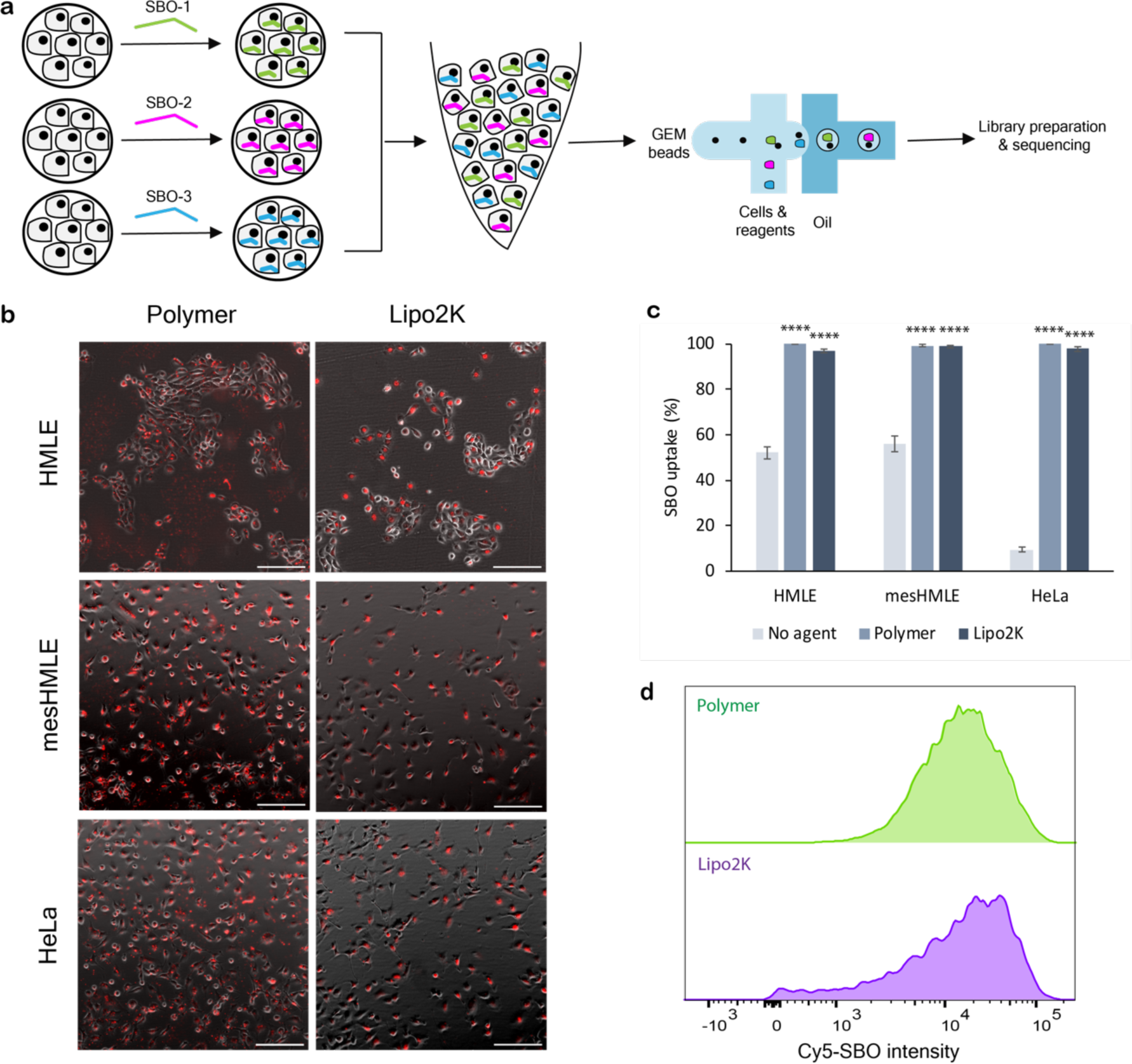
Polymer transfection results in high, uniform SBO delivery for scTECH-seq. **(a)** Schematic of scTECH-seq workflow. Coloured lines represent specific SBOs. Samples were transfected with SBOs, which were then collected, pooled together and processed for single cell RNA sequencing using 10x Genomics Chromium controller. **(b)** Fluorescent microscopy images showing SBO (red) uptake in cells transfected with polymer and Lipofectamine (Lipo2K) in three cell lines; HeLa, HMLE and mesHMLE. Scale bar 100 μm. **(c)** Mean percentage of SBO+ cells for HeLa, HMLE, and mesHMLE cells (two biological replicates, *n* = 6). Significance calculated by t-test, *p* < 0.0001. **(d)** The uniform distribution intensity from the Cy5-conjugated SBO indicates a uniform distribution of SBO in the polymer transfected cells (two biological replicates, *n* = 6).

We compared efficiency of cellular barcoding between our scTECH-seq method and lipid-based oligonucleotide delivery systems, which has been used previously in cell hashing approaches^18^. For scTECH-seq, we used polyplex-based delivery using our previously reported dendritic polymeric system as SBO delivery agents^19^. Our polymer fully bound the SBOs and formed polyplexes ∼100 nm in diameter with a positive surface charge, that were not toxic to cells over 24 h (Supplementary Fig. S1). We initially validated barcoding with Cy5-labelled SBOs using flow cytometry and fluorescence microscopy in three cell lines; HeLa, immortalised human mammary epithelial cells (HMLE) and human mammary epithelial cells that had transitioned to a stable mesenchymal phenotype following TGF-β treatment (mesHMLE) (Fig. 1b-d). Lipofectamine 2000 (Lipo2K) and the polyplexes both demonstrated near-quantitative barcoding efficiency (Supplementary Fig. S2).

Importantly we observed that cells barcoded via polymer-based delivery had a more uniform, narrow uptake of the SBOs in comparison to Lipo2K, indicating that each cell barcoded via polymer-based delivery has similar number of barcodes per cell (Fig. 1d). This is critical in cell hashing technologies as broad distributions of barcodes per cell can lead to a greater possibility of losing cells due to thresholding during the demultiplexing steps. Indeed, we found that the rate of recovery of cells using scTECH-seq is over 72% when compared to lipofectamine-based barcoding, which we found to be approximately 44% (Supplementary Fig. S3). Thus, scTECH-seq has the potential to provide greater scalability and resolution for multi-time point or multi-cell type experiments.

### scTECH-seq reveals multiple unsynchronised metastable states in EMT

We performed scTECH-seq on HMLE incubated with 2 ng/ml TGF-β1 over a twelve day time course, and collected samples at 0, 4, 8, and 12 days, as well as stable mesHMLE cells (Fig. 2a). These timepoints were selected based on immunofluorescence for the epithelial marker, E-cadherin, and mesenchymal marker, vimentin, along with qPCR analysis, which demonstrated an overall reduction in E-cadherin and an increase in vimentin levels consistent with previous EMT observations^20^ (Supplementary Fig. S4). For single-cell profiling, samples were multiplexed via scTECH-seq and captured on the 10x Genomics Chromium controller using the 5′ Gene Expression kit. In total 26,247 single cells were sequenced in 4 channels, with a median of 5,061 genes detected per cell. Upon demultiplexing the cells using the conventional Seurat method, a significant proportion of our data was unable to be assigned a unique single SBO, resulting in lower heterogeneity resolution (Supplementary Fig. S5). Therefore, to demultiplex the cells, a novel deconvolution algorithm called D-score was developed for SBO data. Existing methods (*e.g.,* Seurat HTODemux) use a global read threshold for each barcode to distinguish between background and signal, and assumes that each barcode has equally high reads across all cells. D-score instead calculates a polarisation score for each barcode based on the local reads per cell, such that a high read count from a single barcode from a single cell also results in a high score for that particular barcode, and conversely reads for many barcodes from the same cell result in low scores for each barcode. D-score then assigns a single barcode to a cell. Alternatively, a cell is given an ‘unassigned’ status if there is ambiguity.

**Fig. 2.**
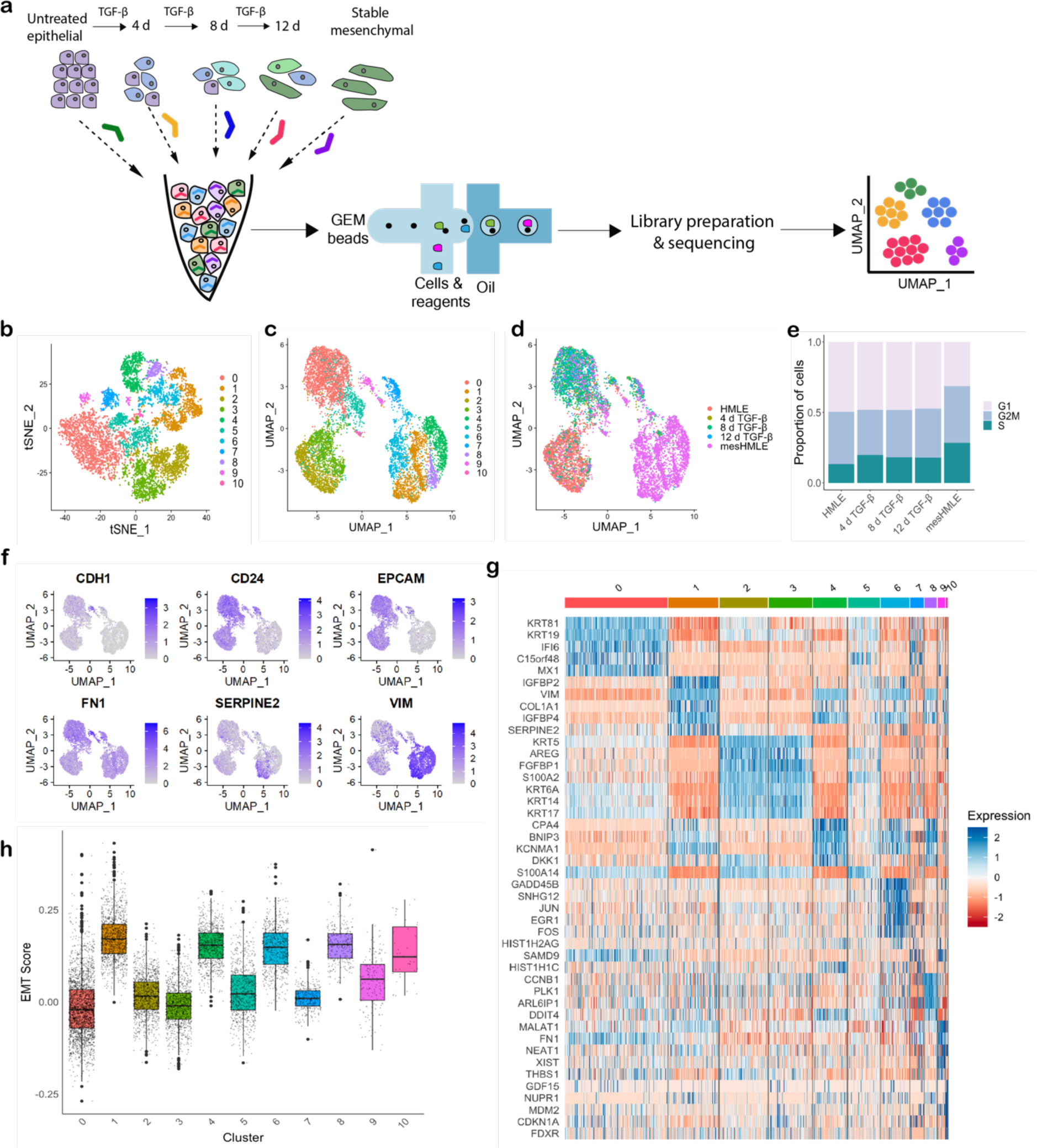
Single cell RNA-sequencing of TGF-β induced EMT in human mammary epithelial cells (HMLE) **(a)** Schematic representation of the samples collected for scTECH-seq, library preparation and data analysis. Cells collected for sequencing include untreated HMLEs, HMLE cells after 4, 8, and 12 days treatment with 2 ng/ml TGF-β1, and mesHMLEs. Coloured lines represent sample specific SBOs. **(b)** t-SNE representation of the cell sub populations. Colours represent different cell clusters. **(c)** UMAP visualisation of the TGF-β EMT timecourse cells. Colours represent different cell clusters. **(d)** UMAP visualisation of the TGF-β EMT timecourse, with colours indicating different samples. **(e)** TGF-β induced EMT does not cause cell cycle arrest. Stable mesHMLE cells showed a high proportion of cells in S phase. **(f)** Log-normalised gene expression of epithelial (*CDH1, CD24, EPCAM*) and mesenchymal (*FN1, SERPINE2, VIM*) markers. **(g)** Top five differentially expressed genes in each cluster (*p* < 0.05). **(h)** Boxplots of cell cluster EMT scores, determined by singscore. Black lines indicate first quartile, median and third quartile EMT score. Individual cell scores are shown as grey dots. Outliers are displayed as large solid black dots.

The Seurat R package was used to employ unsupervised t-distributed stochastic neighbour embedding (tSNE) and uniform manifold approximation and projection (UMAP) clustering (Fig. 2b-d). Given that the TGF-β signalling pathway can induce cell cycle arrest via the upregulation of cyclin-dependent kinase inhibitor p21^21,22^, we used canonical biomarkers to check that cells within samples were not arrested (Fig. 2e). We identified eleven main cell states, two made up of 90% (cluster 2) and 83% (cluster 3) untreated HMLE cells, and four populations arising primarily from mesHMLE cells (89.6%, 93.4%, 89.4%, 93.7%; clusters 1, 4, 6, 8 respectively) (Fig. 2c-d). Expression of canonical EMT gene markers indicated epithelial, mesenchymal and metastable hybrid E/M cell populations within the dataset (Fig. 2f). A main cluster of 2,326 cells (cluster 0), primarily consisting of cells from the samples treated with TGF-β, displayed moderate expression levels of both epithelial and mesenchymal gene markers indicating a potential E/M hybrid population (Fig. 2c-d). We identified 6,181 differentially expressed genes (DEGs) upregulated between the eleven cell clusters and a mean 268 DEGs for each sample, using log_2_ fold change threshold 0.25 and *p*-value < 0.05 (Fig. 2g). The top 5 DEGs for cluster 0 have low expression across all other cell subsets, indicating markers specific to hybrid TGF-β EMT cells (Fig. 2g). We next used Gene Set Enrichment Analysis (GSEA) to visualise pathway enrichment. We observed a positive correlation between EMT and TGF-β signalling pathways, validating TGF-β as an effective EMT trigger in this cell model (Supplementary Fig. S6). Furthermore, we observed enrichment in TGF-β, Wnt, Notch and MAPK pathways in untreated and TGF-β stimulated cells (Supplementary Fig. S6). TGF-β triggers crosstalk interactions with Notch and Wnt/β-catenin signalling pathways^23–26^ resulting in cell differentiation, tumorigenesis and acquisition of migratory potential^23,24,27^. Hence, high expression of these pathways are indicative of cancer stem cell populations. Clusters consisting predominantly of stable mesenchymal cells exhibited low Notch, ERK1/ERK2, p38/MAPK and JNK pathway enrichment (Supplementary Fig. S6), suggesting that once cells complete EMT and fully adopt mesenchymal status there is no need to maintain high levels of transitional signalling.

Next, we assigned each cell subset an EMT score using the singscore R package^28,29^ (Fig. 2h). Singscore uses a rank-based statistic scoring system to analyse gene expression profiles and compare them to up- and down-regulated TGF-β EMT gene signatures. An EMT score of zero is indicative of an epithelial cell state and increasing positive scores correspond to an upregulation of genes associated with a mesenchymal profile. Since single cells from each sample belonged to various clusters, EMT scores were calculated for each subpopulation identified in the tSNE and UMAP clustering. Cell populations that displayed both epithelial and mesenchymal markers also demonstrated transitional EMT scores, indicating multiple EMT hybrid states. In addition, cells within a given cluster showed a range of scores, highlighting the difficulty of counting a definite number of hybrid E/M states. This further demonstrates a non-linear dynamic transition during EMT.

### Metastable clusters are unsynchronised in transitions

The top 1,000 DEGs from the mesHMLE cells were selected and the Monocle v2 R package^30–32^ was used to construct a TGF-β EMT trajectory and order cells accordingly, given that cells do not progress through EMT synchronously. The trajectory showed that most cells from 4, 8, and 12 d TGF-β stimulation did not reach full mesenchymal state (*i.e.,* trajectory endpoint), rather cells from these samples clustered at the epithelial-like end of the construct (*i.e.,* zero pseudotime) (Fig. 3a-c). We also confirmed the decrease of common epithelial markers and subsequent increase in expression of mesenchymal genes over pseudotime (Supplementary Fig. S7). *CDH1* expression decreased with TGF-β treatment while *VIM* increased, consistent with the immunofluorescence results. *CD24* and *EPCAM* expression also decreased with TGF-β treatment as anticipated, and similarly, mesenchymal markers *VIM* and *SERPINE2* expression were highly upregulated in E/M hybrid clusters and mesenchymal cells. Differential gene expression (qval < 0.0001) altered with pseudotime, displaying downregulation of epithelial genes (Fig. 3c). Notably, genes in the S100 family were significantly altered with pseudotime. The S100 family members are known to be involved in biological processes including cell cycle progression, differentiation and disease development^33,34^. *S100A6*, encoding for calcium-binding protein A6, promotes EMT via the Wnt/Beta-catenin signalling pathway and increased in expression with pseudotime. *S100A11*, *S100A2* and *S100A14* genes are associated with tumour invasion and metastasis^35–38^. Expression of these genes varied during EMT, displaying high expression levels within the middle of the trajectory, indicating that cells at these points may possess cancer stem cell traits.

**Fig. 3.**
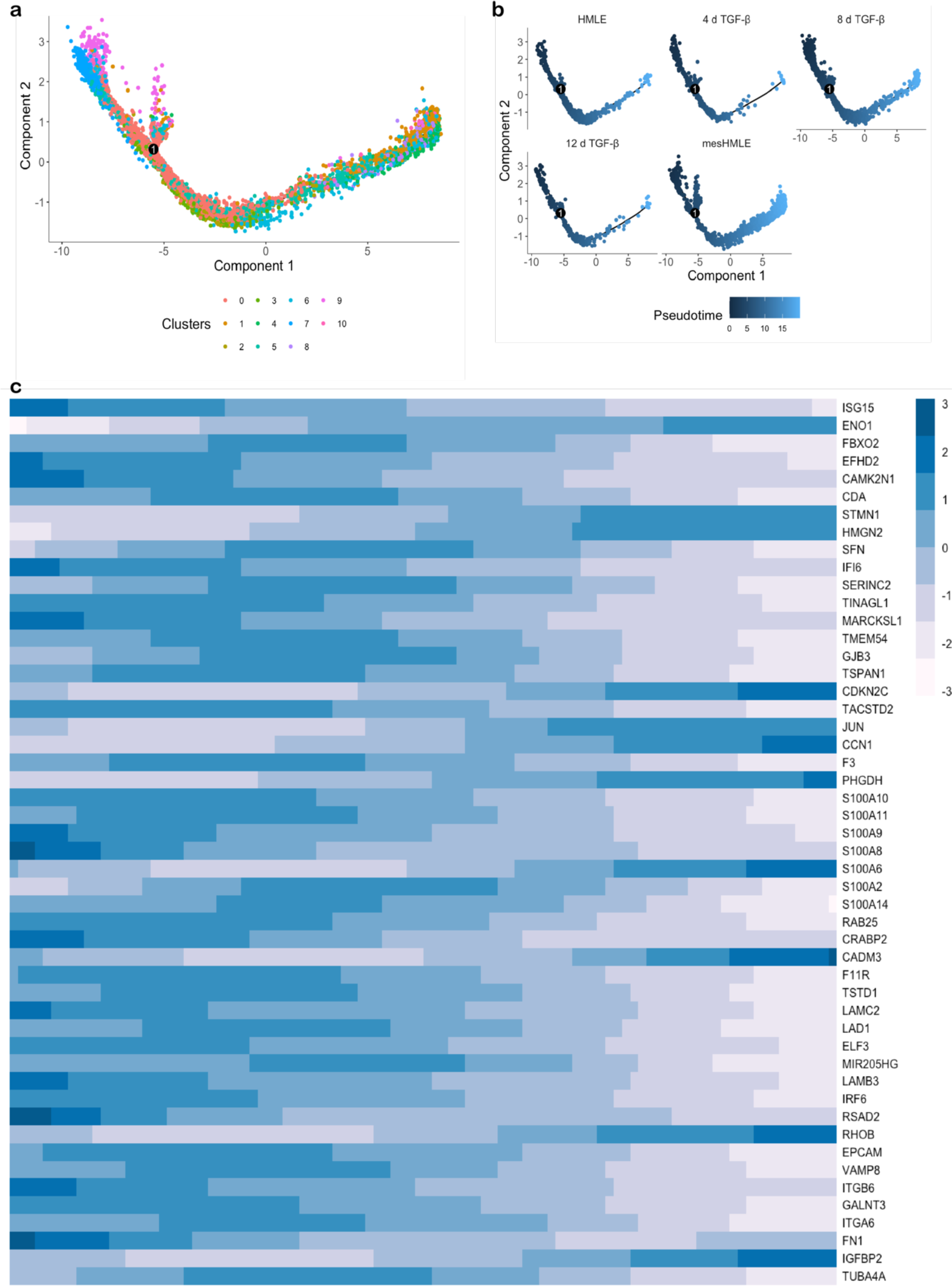
Trajectory analysis of TGF-β induced EMT. **(a)** Trajectory of TGF-β EMT, coloured by cluster. A single branch point in the trajectory is indicated by the number 1 in a black circle. **(b)** Pseudotime analysis showing the distribution of cells based on sample. **(c)** Expression of the top 50 differentially expressed genes (qval < 0.0001) with pseudotime.

### G4 structures regulate transcription of EMT genes

Noncanonical secondary DNA structures called G-quadruplexes (G4s) have been identified as dynamic regulators of transcription, replication, and genome stability. Promoter regions are highly enriched in sequences that fold into G4 structures, where their formation acts to alter gene expression^39–42^. G4 structures have been identified as important molecular targets for regulating the expression of oncogenes that have undruggable gene products.

We used G4-Grinder prediction^43^ to identify which TGF-β EMT DEGs contain G4 sequences in their promoters that may be implicated in the regulation of gene expression. G4-seq, a polymerase-stop based assay, revealed >700,000 G4-forming structures in the human genome^44^. Further, chromatin immunoprecipitation sequencing has revealed approximately 10,000 G4 structures in human chromatin, mostly within regulatory and nucleosome-depleted regions^45^. We identified that of the 6,181 DEGs in TGF-β EMT (p < 0.05, log threshold 0.25), more than one third of these genes (2,250/6,181) contain one or more predicted G4 sequences (pG4) in their promoter (Fig. 4a, Supplementary Table S1). Further analysis of genes containing promoter pG4s revealed that these genes are involved in cell morphogenesis and mitotic cell cycle (Fig. 4b) and are upregulated in cancer pathways (Fig. 4c). Given this, we employed scTECH-seq to examine gene expression profiles of HMLE cells in the presence and absence of theG4-stabilising ligand pyridostatin (PDS)^46^. Under G4 stabilisation conditions we saw an increase in expression of epithelial markers *KRT19* and *CDH1,* both of which are normally silenced with EMT thereby contributing to increased cell proliferation and migration (Fig. 4d). Genes associated with EMT and a mesenchymal phenotype such as *VIM* and *EGR1* were silenced with PDS treatment, while others (eg. *KRT81*, *IGFBP2*, *NEAT1*) were upregulated. Samples treated with PDS achieved a negative EMT score by the singscore algorithm, indicating an overall upregulation in epithelial genes and/or downregulation of mesenchymal genes (Fig. 4e).

**Fig. 4.**
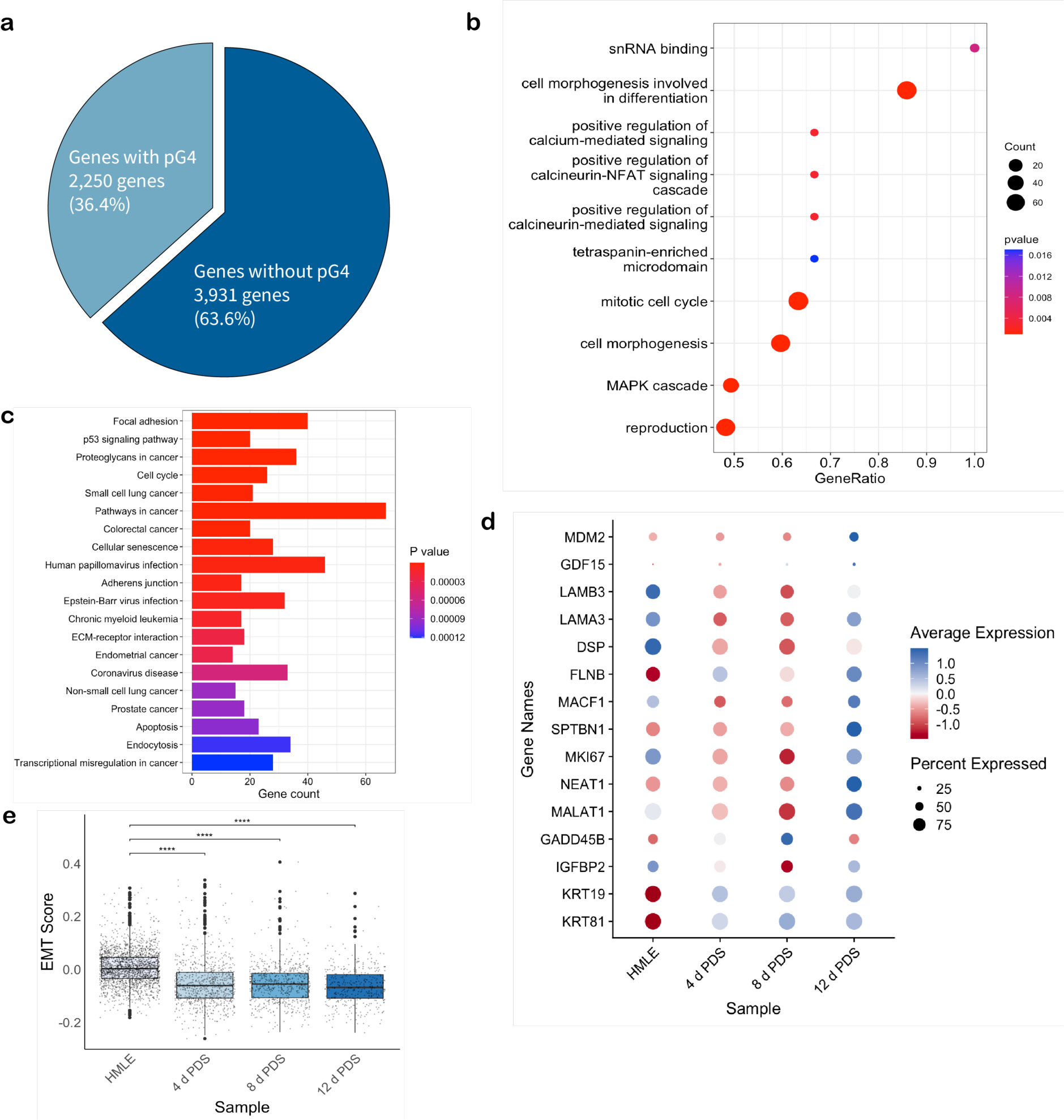
Differentially expressed genes from TGF-β induced EMT that contain putative G4 sequences in gene promoter regions are important for cell regulation. **(a)** Visual representation showing that 36.4% of differentially expressed genes in TGF-β induced EMT contain predicted G4 sequences (pG4) within the promoter regions. **(b)** Gene ontology analysis of the genes that contain pG4, showing pathways associated with cell growth and development are upregulated. **(c)** KEGG pathway analysis of the genes in EMT with pG4 show that G4 structures play a role in regulating cancer progression. **(d)** TGF-β EMT genes with predicted G4s in promoter in HMLE cells treated with PDS for 4, 8, and 12 days alter in expression after G4 stabilisation. **(e)** HMLE cells treated with PDS display an epithelial-like EMT score, calculated using singscore, indicating that genes upregulated with TGF-β EMT are downregulated with PDS treatment. Box plot shows minimum, first quartile, median, third quartile and maximum EMT score. \*\*\**p* < 0.001.

We next examined G4 formation throughout EMT using confocal microscopy. In addition, we examined i-motif (iM) formation, secondary structures that form in cytosine-rich sequences that have an interdependent relationship with G4 structures^47^. We used structure-specific antibodies (BG4^48^ and iMab^49^) to detect and quantify G4 and iM structures respectively, at different stages of EMT (Fig. 5a). E-cadherin and vimentin immunofluorescence were used to monitor EMT progression as epithelial and mesenchymal markers, respectively (Fig. 5a). G4 structure formation was most prominent in epithelial cells and decreased overall as EMT progressed (Fig. 5b). Conversely, iM formation was significantly higher during EMT progression and in the stable mesenchymal cells compared to the original epithelial population (Fig. 5c). We noted that G4 and iM quantification showed large variations at intermediate time point (four, eight and twelve days TGF-β). This agreed with our findings using scTECH-seq that metastable intermediates are multivariant in nature. We further confirmed similar trends in G4 and iM quantification in the PMC42 breast cancer cell model, where the epithelial phenotype (named PMC42-LA) was spontaneously derived via mesenchymal to epithelial transition from its mesenchymal counterpart, PMC42-ET^50^. iMs were more abundant in PMC42-ET cells compared to the epithelial PMC42-LA, which showed a higher number of G4 structures, and generally corresponded to the EMT states of the clonal populations derived from individual PMC42-LA cells (Supplementary Fig. S8). Since G4 and in particular iM stability are impacted by pH changes, we measured differences in intracellular pH during EMT. We found that while intracellular pH did fluctuate in TGF-β induced EMT, there was little change between HMLE cells and mesHMLE cells (Supplementary Table S2).

**Fig. 5.**
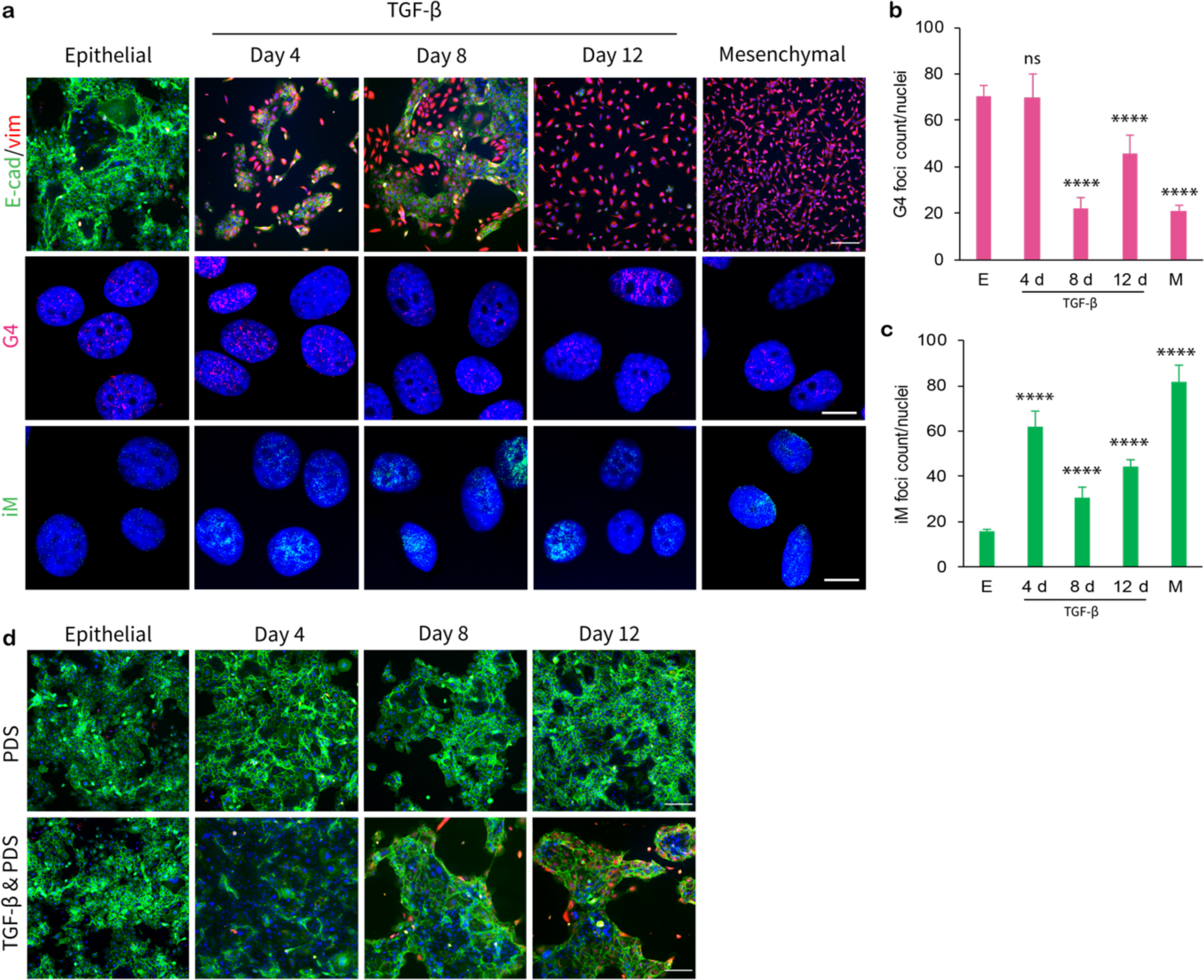
Formation of secondary DNA structures G4s and iMs alter with different EMT phenotypes. **(a)** E-cadherin (green) and vimentin (red) immunofluorescence, scale bar 200 μm. G4 (pink) and i-motif (green) immunofluorescence in cell nuclei (blue), scale bar 20 μm. **(b)** G4 and **(c)** iM quantification, showing mean and SEM (*n* = 67–119). Statistical significance is shown relative to epithelial (E) foci count. **(d)** HMLE cells treated with 0.4 μM PDS in the absence and presence of 2 ng/mL TGF-β display high levels of epithelial marker E-cadherin (green) and low levels of mesenchymal protein vimentin (red) by immunofluorescence (scale bar 200 μm). **** p < 0.0001, ns: p > 0.05.

Given that the HMLE cells displayed higher G4 formation compared to the mesHMLE cells, we next investigated the impact of G4 stabilisation in the presence and absence of EMT trigger TGF-β. To stabilise G4 structures cells were treated with 0.4 μM pyridostatin (PDS) and G4 formation was visualised with BG4 immunofluorescence (Supplementary Fig. S9). G4 stabilisation with PDS in the absence of TGF-β increased the number of G4 structures at the earlier timepoints, 4 and 8 days, when compared to TGF-β treated cells, with no significant alterations observed at day 12. G4 stabilisation at all timepoints in the presence and absence of the EMT trigger TGF-β, resulted in low vimentin protein levels while maintaining consistent E-cadherin protein cell surface expression (Fig. 5d), indicating a relationship between G4 formation and these EMT markers, where G4 stabilisation favours the epithelial phenotype. Collectively, the change in EMT markers observed with G4 stabilisation prompted us to investigate the transcriptomic implications of G4 stabilisation during TGF-β-induced EMT.

### scTECH-seq of G4 stabilised EMT

HMLE cells treated with 0.4 μM PDS, in the presence or absence of TGF-β for four, eight or twelve days were multiplexed and sequenced at the single cell level. Unsupervised clustering showed ten subpopulations (Fig. 6a-c). The PDS treatment alongside TGF-β stimulation resulted in a scattering of cells across various clusters, whereas untreated HMLE cells belonged primarily to clusters 2 and 3 (Fig. 6d). Given that G4 formation is cell cycle dependent, being most prominent during S phase of the cell cycle, we first confirmed that PDS treatment did not induce cell cycle arrest (Fig. 6e). TGF-β + PDS treated cells dominated clusters that displayed an increase in expression of genes with G4 structures (Fig. 6f). Given that G4 structures have been implicated indirectly in chromatin remodelling, we were not surprised to see differential expression of Histone encoding genes (Fig. 6f). We observed that cluster 8, a small cluster (3.37% of the population), displayed mesenchymal characteristics including high expression of *VIM* and *IGFBP4* and low *CDH1* and *EPCAM* expression (Fig. 6f-g). This suggests that the combined treatment of TGF-β and PDS resulted in only a small population of cells to transition into a mesenchymal-like phenotype. The high expression of epithelial markers observed with PDS treatment indicate that G4 stabilisation favours an epithelial-like state. EMT scoring via the singscore R package^28,29^ revealed that the majority of cells exhibit an epithelial-like profile, with only one cluster (cluster 8) exhibiting a high mesenchymal score (Fig. 6h). A negative EMT score indicates genes that are upregulated with canonical TGF-β EMT (i.e., mesenchymal genes) are in fact downregulated, whereas genes that are expected to decrease with EMT (i.e., epithelial genes) are still being expressed. This suggests that clusters with EMT scores below zero have upregulated epithelial gene expression profiles. Clusters consisting of cells that were treated with both TGF-β and PDS showed negative EMT scores, supporting the trajectory findings that G4 stabilisation is altering the expression of important EMT genes, thereby impacting overall EMT progression. At all timepoints, PDS treatment resulted in a lower EMT enrichment score compared to the respective TGF-β counterparts (Supplementary Fig. S10), providing further evidence that G4 stabilisation via PDS impacts the expression of EMT genes.

**Fig. 6.**
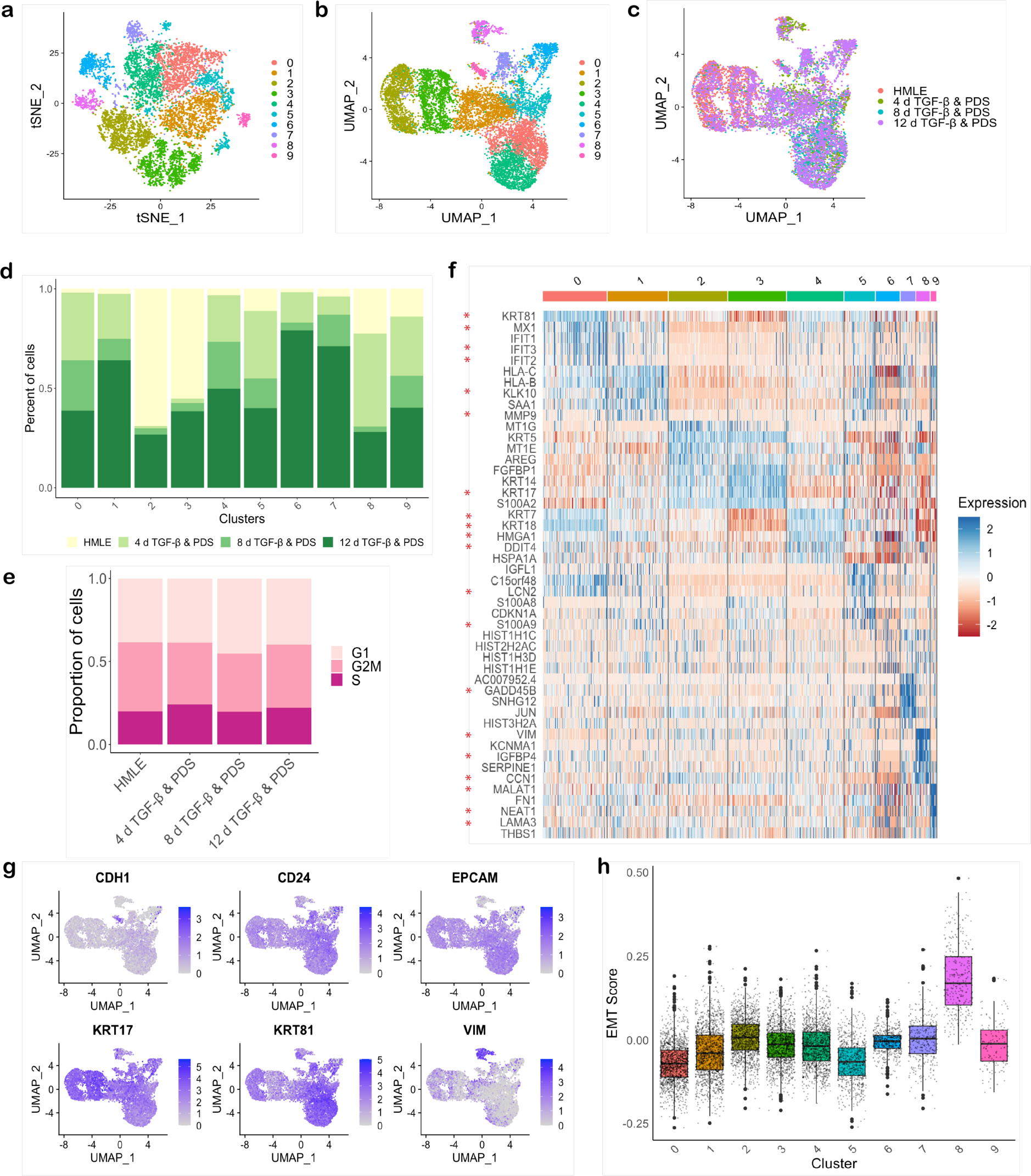
Ligand-induced G4 stabilisation alters TGF-β EMT. **(a)** tSNE and **(b)** UMAP visualisation of 23,032 cells analysed by scTECH-seq. **(c)** UMAP visualisation of cells annotated for the sample of origin. **(d)** Distribution of samples across subpopulation clusters identified from tSNE and UMAP clustering. **(e)** Proportion of cells from each sample at cell cycle phases. **(f)** Differential expression analysis (*p* < 0.05, log_2_FC < 0.25) was performed comparing clusters identified in the tSNE and UMAP. Genes with red asterisks contain predicted G4 sequences. **(g)** The expression of known EMT markers. **(h)** Boxplots of EMT scores, determined by singscore, for cell clusters reveal that the majority of subpopulations display epithelial-like phenotypes. Black lines indicate first quartile, median and third quartile EMT score. Individual cell scores are shown as grey dots. Outliers are displayed as large solid black dots.

G4 Grinder analysis showed that more than one third of genes (111/302 genes) used to calculate EMT scores contain pG4 in promoters, including key EMT genes *CDH1*, *ESRP1, VIM* and *TGF-B1* (Supplementary Table S3 & S4). Pseudotime trajectory analysis for cells treated with both TGF-β and PDS revealed seven cell fates (labelled A-G) (Fig. 7a-b), with gene expression patterns differing at opposite ends of the trajectory (Supplementary Fig. S11). The untreated HMLE cells were placed towards the middle of the trajectory, rather than at the start point at pseudotime 0 (ie. cells that display the most dissimilar gene profiles to the mesHMLEs) which primarily consisted of cells that underwent TGF-β and PDS treatment. This observation indicates that targeting G4 formation can be used to alter EMT progression. Lastly, we compared the expression of DEGs from our TGF-β dataset in PDS treated cells (Fig. 7c). Genes associated with cell proliferation, invasion and metastasis, including *SNHG12*^51^, *XIST*^52^ and *FN1*^49^, decreased in expression with PDS. *SMAD9*, a protein involved in TGF-β signalling, and *THBS1*, a TGF-β target gene, displayed lower expression levels at 4, 8 and 12 d timepoints with PDS treatment. Indeed, PDS treatment resulted in a shift in pathway and gene ontology enrichment (Supplementary Fig. S12), suggesting that targeting G4 formation leads to downstream impacts on EMT signalling pathways.

**Fig. 7.**
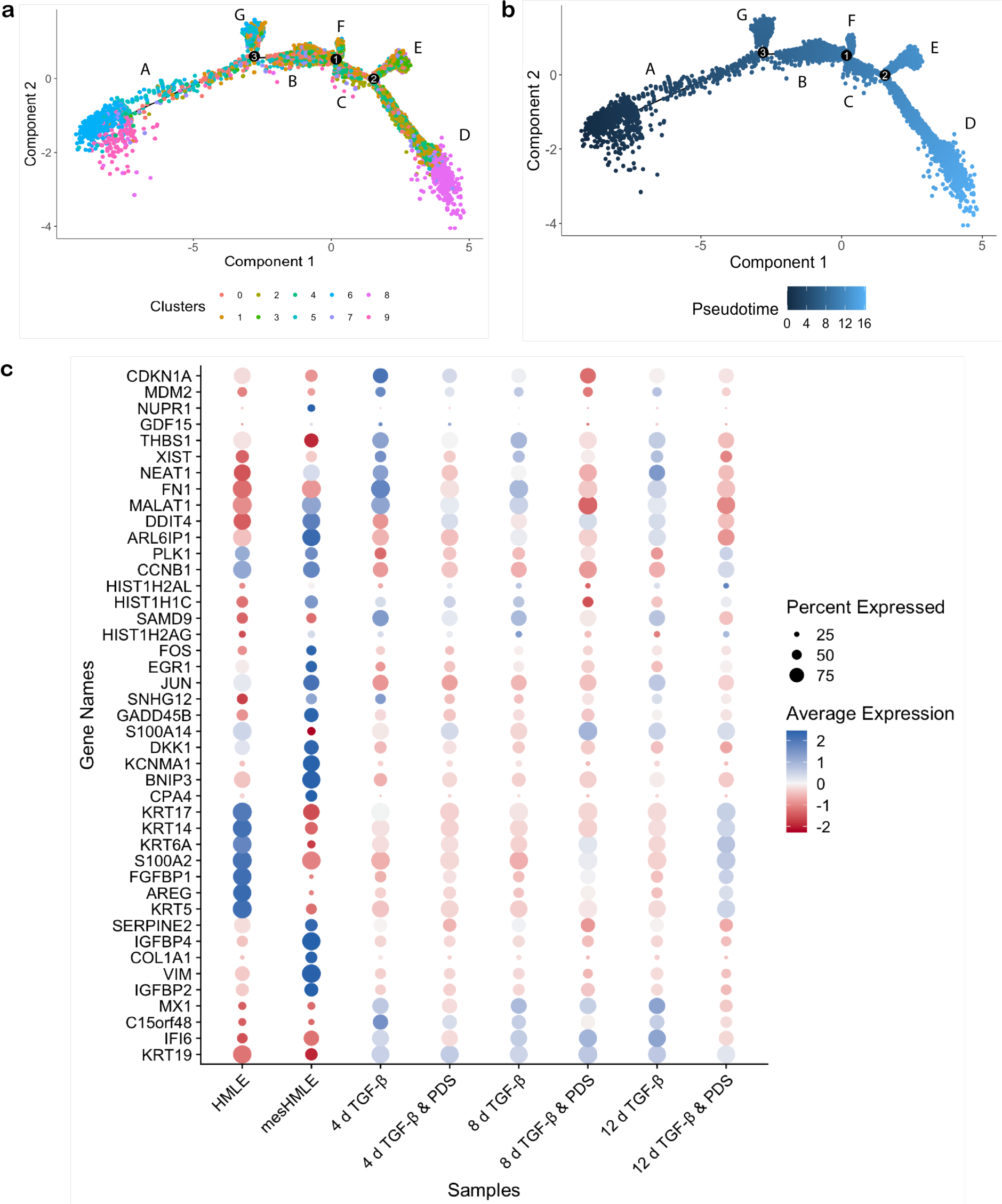
G4 stabilisation via PDS treatment alters TGF-β triggered EMT. **(a)** Trajectory analysis calculated via Monocle2. Multiple nodes along the trajectory indicate that TGF-β + PDS treatment causes multiple cell fates, indicated by numbers. Seven cell states (A-G) were identified. Cells are annotated based on cell cluster assignment from UMAP and tSNE clustering. **(b)** Pseudotime trajectory analysis of cells treated with TGF-β + PDS. **(c)** Addition of PDS alters the expression of gene markers identified from TGF-β triggered EMT (Fig. 1).

## Discussion

Herein, we have described scTECH-seq for multiplexing samples for single cell RNA-sequencing. Multiplexed methods have proven essential for removing batch effects and significantly reducing cost of sequencing studies. scTECH-seq utilises our in-house polymer to deliver DNA barcodes (SBOs) into target cells with both high transfection efficiency and a more uniform uptake of SBOs in comparison to the commercially available transfection agent Lipofectamine 2000. scTECH-seq is an antigen-independent multiplex method that labels every cell within a given sample and is compatible with low cell numbers thus making it suitable for multiplexing samples with rare cells. Our method offers the unique advantage of multiplexing samples for 5’ gene expression, with current protocols such as 10x Genomics’ Cell Plex only suitable for 3’ gene expression profiling.

The application of scTECH-seq enabled the identification of multiple metastable, hybrid populations in TGF-β induced EMT. Pseudotime analysis revealed the dynamic nature of this biological transition, with cells treated with TGF-β for 4, 8, and 12 days distributed along the EMT spectrum, highlighting that EMT does not occur synchronously. Despite the highly dynamic and complex nature of this process, we demonstrate how G4 DNA structures are involved in regulating EMT. By targeting G4 structures, we showed we could alter EMT progression at the gene level and in pathway signalling.

## Methods

### Polymer characterisation

The dendritic polymer was synthesised as outlined by Kretzmann *et al.*^19^ using a 25% GMA backbone and generation 5.0 poly(amido amine) dendrons. Polymer-SBO binding was assessed by a gel retardation assay. Polymer solutions were mixed with 1 μg SBO at different amine-to-phosphate (N/P) ratios and incubated at 25 °C for 30 min. Samples were electrophoresed on a 1% w/v agarose gel in sodium borate buffer at 120 V for 80 min. For dynamic light scattering (DLS) and zeta potential measurements, various N/P polymer ratios were mixed with SBO (1 μg) and incubated for 30 min. Solutions were diluted to 400 μL and measurements were taken in triplicate. Reported values are the mean ± SD for average peak size and zeta potential.

### Cell culture

Immortalised human mammary luminal epithelial cells (HMLE) were cultured in HUMEC media (Gibco, cat# 12753-018). Stable mesenchymal HMLE cells (mesHMLE) were maintained in Weinberg media (DMEM/F12 1:1 supplemented with 5% v/v FBS, 10 μg/mL insulin, 20 ng/mL EGF and 0.5 µg/mL hydrocortisone)^7^. Cell lines were maintained at 37 °C in a humidified incubator with 5% CO_2_ and passaged at 70–80% confluency using 1× TrypLE Express. To study EMT, HMLE cells were cultured in Weinberg media supplemented with 2 ng/mL TGF-β1 for 0–12 d. In G4 stabilisation experiments, Weinberg media was supplemented with 0.4 μM PDS or 2 ng/mL TGF-β1 and 0.4 μM PDS. For immunofluorescence studies, cells were seeded onto sterile glass coverslips at 200,000 cells/well 24 h prior to fixation.

### Transfection protocol

Cells (HeLa, HMLE and mesHMLE cells) were seeded at 160,000 cells/well in a 24-well plate and left overnight at 37 °C and 5% CO_2_. Polymer solutions and SBO were diluted to working concentrations in Opti-MEM (Gibco cat# 31985070). SBO was diluted to give 23 pmol in 35 μL Opti-MEM. Polymer and SBO solutions were mixed gently but thoroughly and incubated for 25 min. Cells were washed with 1×PBS once to remove serum and media was replaced with 150 μL Opti-MEM. 65 μL Polymer-SBO complexes were added and cells were incubated at 37 °C for 4 h. Lipofectamine 2000 was used according to the manufacturer’s protocol under the same incubation conditions. Transfection efficiency was visualised with an epi- fluorescence microscope. For quantification via flow cytometry, cells were incubated with 250 μL 1× TrypLE for 10 min and collected with 250 μL 2×FACS buffer (6% v/v FBS, 2 mM EDTA in 1×PBS).

### Cell toxicity assay

HMLE cells were plated at 3,000 cells/well in a 96-well plate in 100 μL HuMEC media and incubated overnight. The next day, media was replaced with Weinberg media containing PDS, ranging from 0.1–100 μM for 72 h. After 72 h, 20 μL MTS was added to each well and cells were incubated for a further 2 h before absorbance was measured on plate reader at 490 nm.

### scTECH-seq sample preparation

HMLE cells were grown in Weinberg media supplemented with TGF-β, TGF-β and PDS, and just PDS for 4, 8 and 12 days. Media was changed every 2 days. Samples were barcoded, each with a unique SBO as per the transfection protocol above. After 4 h, transfection media was removed and cells were washed twice with PBS. Cells were collected with TrypLE and counted. Cells were resuspended in 0.04% w/v bovine serum albumin (BSA) and pooled together in equal numbers at a final concentration of 1,000 cells/μL.

### Single cell RNA-seq library preparation and sequencing

Cells were loaded onto a 10x Genomics Chromium Controller, Chip K (PN 1000286) and libraries were prepared using the Chromium Next GEM single cell 5’ kit v2 (PN1000263/1000265). Libraries from SBOs were prepared using the 5’ Feature Barcode kit (PN 1000256) following manufacturer guidelines. Illumina iSeq 100 platform was used for initial QC of the libraries and deeper sequencing was conducted on Illumina NovaSeq 6000 in the vendor recommended format) to reach up to 50,000 reads per cell. Fastq files were generated with Illumina bcl2fastq v2.20.0.422, allowing 1 mismatch in each index.

### Single cell data analysis

Transcript alignment and generation of gene count and feature barcode matrices were performed using 10x Genomics Cell Ranger count v6.0.2. Output from four channels was combined either with Cell Ranger aggr (for data QC and initial exploration), or Seurat v4.1.0 in R for all further downstream analysis. A total of 26,247 single cells were sequenced, with a total of 36,601 genes detected and median of 5,061 genes per cell. To demultiplex the cells, a novel deconvolution algorithm called D-score was developed for SBO data. D-score assigns a single hashtag (or ‘unassigned’) to each GEM based on local SBO reads and these were added to the Seurat object metadata. Cells were filtered for 200–80,000 features and less than 10% mitochondrial genes. A total of 23,032 cells were used for downstream analysis and subsetted according to D-score assignment for group analyses. The data was normalised and the top 2,000 variable features were selected for scaling. Principle component, dimension reduction and visualisation (UMAP and tSNE) were performed on the scaled data. The Wilcoxon rank sum test was used to calculate differentially expressed genes (*p* < 0.05 and log threshold 0.25). The Wilcox test was used to calculate statistical significance between samples (*p* < 0.05).

### EMT scoring

Singscore simple scoring was used to calculate EMT scores for clusters and samples^28,29^. To set up the single cell data for singscore, RNA counts were normalised and the RNA matrix was extracted from the Seurat object and made into a summarised experiment. Genes were ranked and scored using the TGF-β EMT up and down gene sets reported in Foroutan *et al.* 2018^28^.

### Trajectory analysis

Monocle v. 2 was used to construct a trajectory in pseudotime^30–32^. The top 1,000 mesHMLE differentially expressed genes were used to plot cells along the trajectory and in pseudotime. The machine learning algorithm plots cells on the learned trajectory based on their differential gene expression profiles. Pseudotime heatmaps show differentially expressed genes (qval < 0.1).

### Enrichment analysis

Single-sample gene-set enrichment analysis were calculated using the GSVA R package (v1.42.0)^54^ using MSigDB hallmark and gene ontology gene sets for EMT and TGF-β signalling pathways. Data normalisation was conducted prior to enrichment. Hallmark and gene ontology biological process sets were downloaded from MSigDB. EnrichR and clusterProfiler R packages were used for KEGG and gene ontology enrichment analysis, using the first 2,000 differentially expressed genes.

### G4 prediction

G4 Grinder^43^ was used to search and analyse potential G4 structures in promoter regions TGF-β EMT genes. Gene promoter sequences were extracted from hg38, 1,000 bp up- and downstream of transcription start site. G4 Grinder default variable settings were used (minimum number of runs = 4), with a minimum acceptable score > 40. Analysis method 3A was selected to detect non-overlapping, size-independent structures.

### BG4 expression and purification

The BG4 pSANG10-3F-scFv plasmid was kindly supplied by Sir Prof. S. Balasumbramanian (University of Cambridge, UK). BL21(DE3) competent cells containing the BG4 plasmid were inoculated overnight at 37 °C, 200 rpm, in ZY medium (10 g tryptone, 5 g yeast extract in 1 L milliQ H_2_O) supplemented with 2% glucose and 50 μg/ml kanamycin. The overnight culture was expanded in autoinduction media (ZY media supplemented with 2 mM MgSO_4_, 0.2× metals mix^55^, 1×5052 (50×5052: 25% w/v glycerol, 2.5% w/v glucose, 10% w/v α-lactose), 1× M (50×M: 1.25 M KH_2_PO_4_, 2.5 M NH_4_Cl, 0.25 M Na_2_SO_4_) and 50 μg/ml kanamycin) at 37 °C, 250 rpm for 6 h. The same culture was incubated overnight at 25 °C, 280 rpm. For BG4 purification, bacterial cultures were centrifuged (30 min, 4000*g* at 4 °C) and resuspended in TES buffer (50 mM Tris-HCl pH 8, 1 mM EDTA, 20% w/v sucrose). TES buffer (diluted 1:5 in milli-Q H_2_O) (supplemented with 10 μL benzonase (Millipore cat# E1014), 2 mM MgSO_4_ and protease inhibitor cocktail (Roche cat# 04693132001)) was added and bacteria was incubated on ice for 15 min before centrifugation for 20 min, 16,000*g* at 4 °C. The supernatant from the bacteria was filtered using a 0.45 μm filter. His-Select nickel affinity beads were equilibrated and evenly distributed among filtered supernatant before incubating at 4 °C for 1 h with rotation. Solutions were centrifuged and supernatant was carefully poured off as to not disrupt beads. Beads were washed with 100 mM NaCl and 10 mM imidazole (pH 8.0) and loaded onto a Ni column. BG4 was eluted with 250 mM imidazole/PBS (pH 8.0) and dialysed overnight at 4 °C. BG4 concentration was determined using Nanodrop and purity was assessed using SDS-PAGE.

### BG4 immunofluorescence

Cells were washed once with 1×PBS + 0.1% v/v Triton-X 100 and 1×PBS. Cells were fixed with 4% paraformaldehyde (PFA) in 1×PBS for 15 min at 25 °C, washed twice with 1×PBS and then permeabilised with 1×PBST for 10 min. Cells were washed with 1×PBS twice followed by blocking with 1×PBS + 2% w/v skim milk powder for 1 h at RT to prevent non-specific binding. Samples were then incubated with 300 nM BG4 antibody diluted in 1×PBS + 1% w/v skim milk powder for 1 h at 37 °C with 5% CO_2_. Next, cells were washed in 1×PBS + 0.1% v/v Tween 20 (1×PBST) followed by incubation with Anti-FLAG (1/1000) (DYKDDDDK tag antibody, #2368, Cell Signal Technology) for 1 h at 37 °C with 5% CO_2_. Cells were washed 3 × 5 min with 1×PBS, followed by incubation with Alexa Fluor 594-conjugated donkey anti-rabbit (1/2000) (#A-21207, Invitrogen) and Hoechst 34580 (1 µg/mL) at 37 °C with 5% CO_2_ for 30 min. Protected from light, samples were washed with 1×PBST and 1×PBS and coverslips were mounted onto clean microscope slides with Fluoromount G (Southern Biotech). Images were acquired with a Nikon A1 RMP confocal using a 60×/1.49 oil immersion objective, and analysed using ImageJ.

### iMab immunofluorescence

Cells were washed once each with 1×PBS + 0.1% v/v Triton-X 100 and 1×PBS. Cells were fixed with 4% PFA in PBS for 15 min at 25 °C, washed twice with 1×PBS and then permeabilised with 1×PBST for 10 min. Cells were washed 2 × 5 min with 1×PBS, followed by overnight blocking with 1×PBS + 2% w/v BSA + 1% w/v skim milk powder at 4 °C. Next, cells were incubated overnight with 200 nM iMab in fresh blocking buffer. Cells were washed 3 × 5 min with 1×PBST and incubated with rabbit anti-FLAG (1/800) for 45 min at 25 °C. After three 1×PBST washes, cells were incubated with Alexa Fluor 594-conjugated donkey anti-rabbit (1/400) and Hoechst 34580 (1 µg/mL) for 45 min at 25 °C. Both anti-FLAG and Alexa Fluor antibodies were diluted in 2% gelatin from cold water fish skin + 0.5% w/v BSA + 0.1% v/v Tween 20 + 1×PBS. Samples were washed twice with 1×PBST and mounted onto clean microscope slides using Fluoromount G (Southern Biotech). Images were acquired with a Nikon A1 RMP confocal using a 60×/1.49 oil immersion objective, and analysed using ImageJ.

### E-cadherin & vimentin immunofluorescence

Incubations were performed at room temperature unless otherwise stated. Cells were fixed with 4% PFA in 1×PBS for 15 min, and washed twice with 1×PBS. Cells were permeabilised with 1×PBST for 10 min at 25 °C, followed by 2 × 5 min washes in 1×PBS. Next, cells were blocked for 1 h in 1×PBS + 2% w/v BSA and then incubated with mouse anti-E-cadherin (1:1000) (Abcam, cat#231303) and chicken anti-vimentin (1:200) (Abcam, cat#24525) diluted in blocking buffer overnight for 18 h at 4 °C. Afterwards, cells were washed 3 × 5 min with 1×PBS + 0.1% v/v Tween 20. Cells were incubated with Alexa Fluor 647-conjugated donkey anti-mouse (1:500) (Abcam, cat#150107), Alexa Fluor 568-conjugated goat anti-chicken (1:500) (Abcam, cat#175477) and Hoechst 34580 (1 µg/mL) for 1 h at 25 °C. Finally, samples were washed twice with 1×PBST and once with 1×PBS before mounting coverslips onto microscope slides with Fluoromount G (Southern Biotech).

### Data and Code availability

The single-cell sequencing datasets generated in this study have been deposited in the National Center for Biotechnology Information Gene Expression Omnibus under accession number GSE225155. The R code for calculating the D-score are available at https://github.com/genomicswa/d-score. The scripts used to generate the plots presented in the main figures are available at https://github.com/Jessica736/scTECH-seq-EMT.

## Acknowledgements

The authors acknowledge Sir Prof. S. Balasubramanian for his gift of the pSANG10-3F-BG4 plasmid and Prof. D.U. Christ for the iMab antibody. We also give thanks to Prof. M. Davis for help in adjusting the singscore R package for single cell data.

This work was funded by the Australian Research Council and National Health and Medical Research Foundation (NHMRC) of Australia. C.W.E. is the recipient of funding from the Government of Western Australia. We would like to acknowledge facilities and assistance at the Centre for Microscopy, Characterisation and Analysis at the University of Western Australia. Library preparation and sequencing was conducted in the Genomics WA Laboratory in Perth, Australia. Genomics WA is supported by BioPlatforms Australia, the State Government of Western Australia, Harry Perkins Institute of Medical Research, Cancer Research Trust, Telethon Kids Institute and the University of Western Australia. We gratefully acknowledge the Australian Cancer Research Foundation and the Centre for Advanced Cancer Genomics for making available Illumina Sequencers for the use of Genomics WA. The Translational Research Institute receives support from the Australian Federal Government.

