## Supplementary Information for "Metastable Intermediates Identified in Epithelial to Mesenchymal Transition are Regulated by G-Quadruplex DNA Structures"

**SUPPLEMENTARY INFORMATION** .......... 3

SUPPLEMENTARY FIGURE 1. POLYMER-SBO COMPLEXES HAVE HIGH TRANSFECTION EFFICIENCY AND LOW CYTOTOXICITY. .......... 3
SUPPLEMENTARY FIGURE 2. SBO DELIVERY INTO CELLS VIA POLYMER TRANSFECTION RESULTS IN HIGH SBO UPTAKE .......... 4
SUPPLEMENTARY FIGURE 3. POLYMER PERFORMS BETTER THAN LIPOFECTAMINE2000 (LIPO2K) AT DELIVERING SBOS FOR MULTIPLEX EXPERIMENTS. .......... 4
SUPPLEMENTARY FIGURE 4. TGF-B STIMULATED EMT IN HMLE CELLS, RESULTING IN THE LOSS OF EPITHELIAL MARKERS AND GAIN OF MESENCHYMAL MARKERS. .......... 5
SUPPLEMENTARY FIGURE 5. SCTECH-SEQ CAPTURES DIVERSITY IN 12-PLEX EMT EXPERIMENT. .......... 6
SUPPLEMENTARY FIGURE 6. EMT SIGNALLING PATHWAYS ARE ENRICHED IN TGF-B INDUCED EMT .......... 7
SUPPLEMENTARY FIGURE 7. GENE EXPRESSION ALTERS WITH EMT PSEUDOTIME. .......... 7
SUPPLEMENTARY FIGURE 8. G4 AND IM IMMUNOFLUORESCENCE IN PMC42 CELL NUCLEI .......... 8
SUPPLEMENTARY TABLE 1. INTRACELLULAR PH (PHI) OF HMLE CELLS CHANGES WITH TGF-B STIMULATION. .......... 8
SUPPLEMENTARY FIGURE 9. PYRIDOSTATIN (PDS) STABILISES G4 STRUCTURES. .......... 9
SUPPLEMENTARY FIGURE 10. PDS LOWERS THE EMT SCORE OF SAMPLES. .......... 10
SUPPLEMENTARY FIGURE 11. TGF-B AND PDS TREATMENT ALTERS GENE EXPRESSION PROFILES. .......... 11
SUPPLEMENTARY FIGURE 12. G4 STABILISATION ALTERS GENE ONTOLOGY IN EMT. .......... 12

**SUPPLEMENTARY METHODS** .......... 13

Supplementary Tables S1, S3, and S4 are located in a separate supplementary spreadsheet.

Supplementary Table S1. G4Grinder analysis of TGF-β differentially expressed genes.
Supplementary Table S3. G4Grinder analysis of singscore's TGF-β down regulated gene list.
Supplementary Table S4. G4Grinder analysis of singscore's TGF-β up regulated gene list.

### Supplementary Information

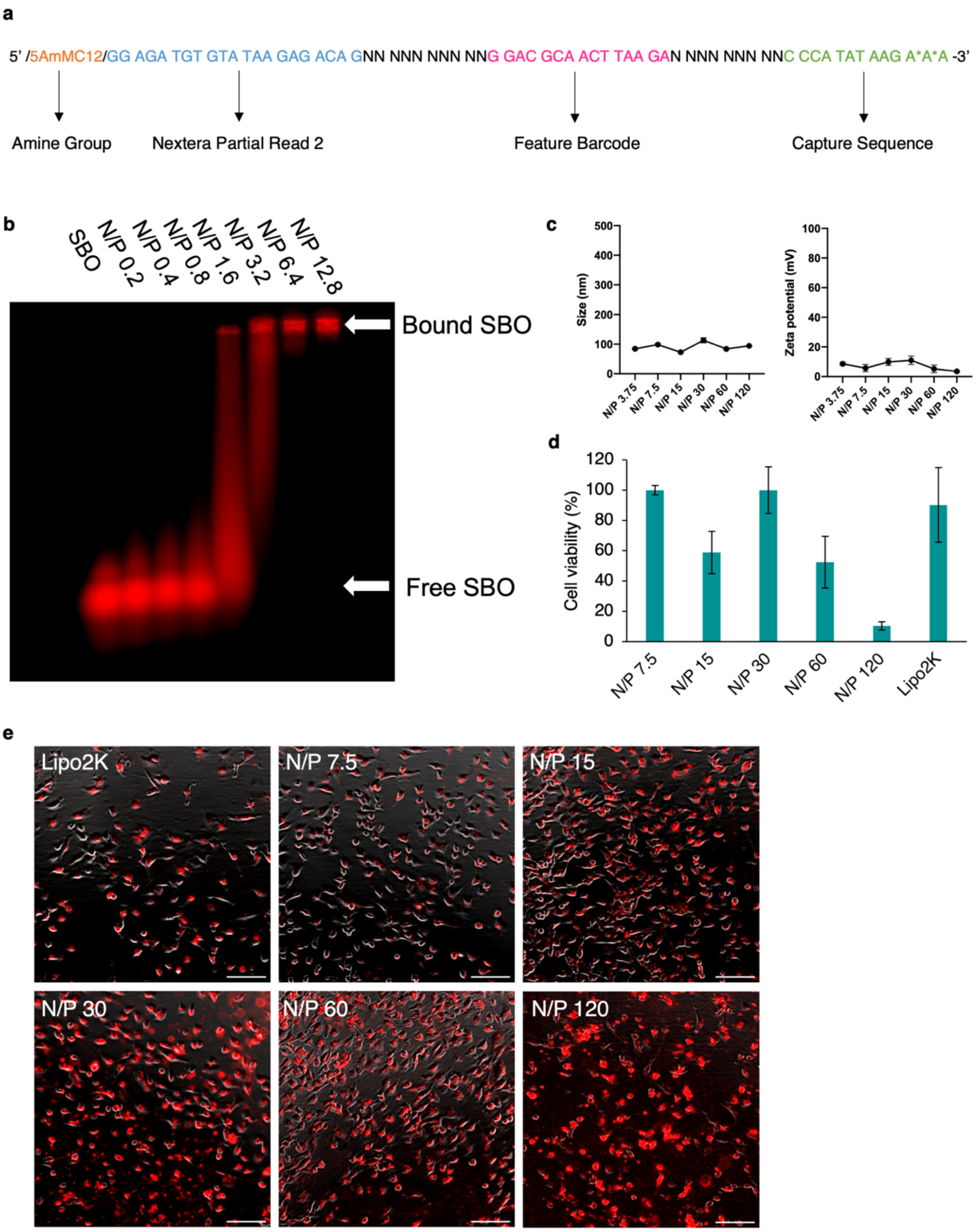


**Supplementary Figure S1. Polymer-SBO complexes have high transfection efficiency and low cytotoxicity. (a)** Sequence structure of SBOs. A SBO contains a 15 nt feature barcode with a 10 nt and 9 nt region of diversity either side (where N = A, T, G, C). The oligonucleotide also contains a 3′ capture sequence (CCCATATAAGA*A*A where * represents phosphorothioate) and a 5′ Nextera partial Read 2 (GGAGATGTGTATAAGAGACAG) with a 5′ $C_{12}$-linked amine group (5AmMC12). **(b)** Gel retardation assay showing that SBO is fully bound to the polymer by amine-to-phosphate ratio (N/P) 6.4. **(c)** Size and surface charge of polymer-SBO complexes. Data is displayed as mean ± S.D. from three measurements. **(d)** Mean cell viability of HeLa cells incubated with polymer-SBO complexes for 48 h. Conditions were conducted in three technical replicates. Data is displayed as mean ± S.D. **(e)** Fluorescence images of polymer and Lipofectamine2000 transfected HeLa cells showing SBO uptake (red) and low toxicity. Scale bar, 500 µm.

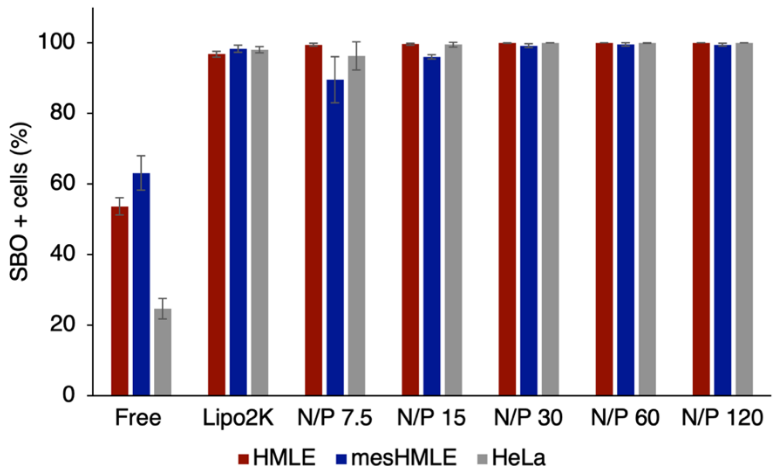


**Supplementary Figure S2. SBO delivery into cells via polymer transfection results in high SBO uptake.** Human mammary epithelial cells (HMLE), mesenchymal mammary cells (mesHMLE) and HeLa cells display > 99% SBO uptake by polymer delivery at N/P 30. Data is displayed as mean ± S.D. calculated from six technical replicates across two biological replicates.

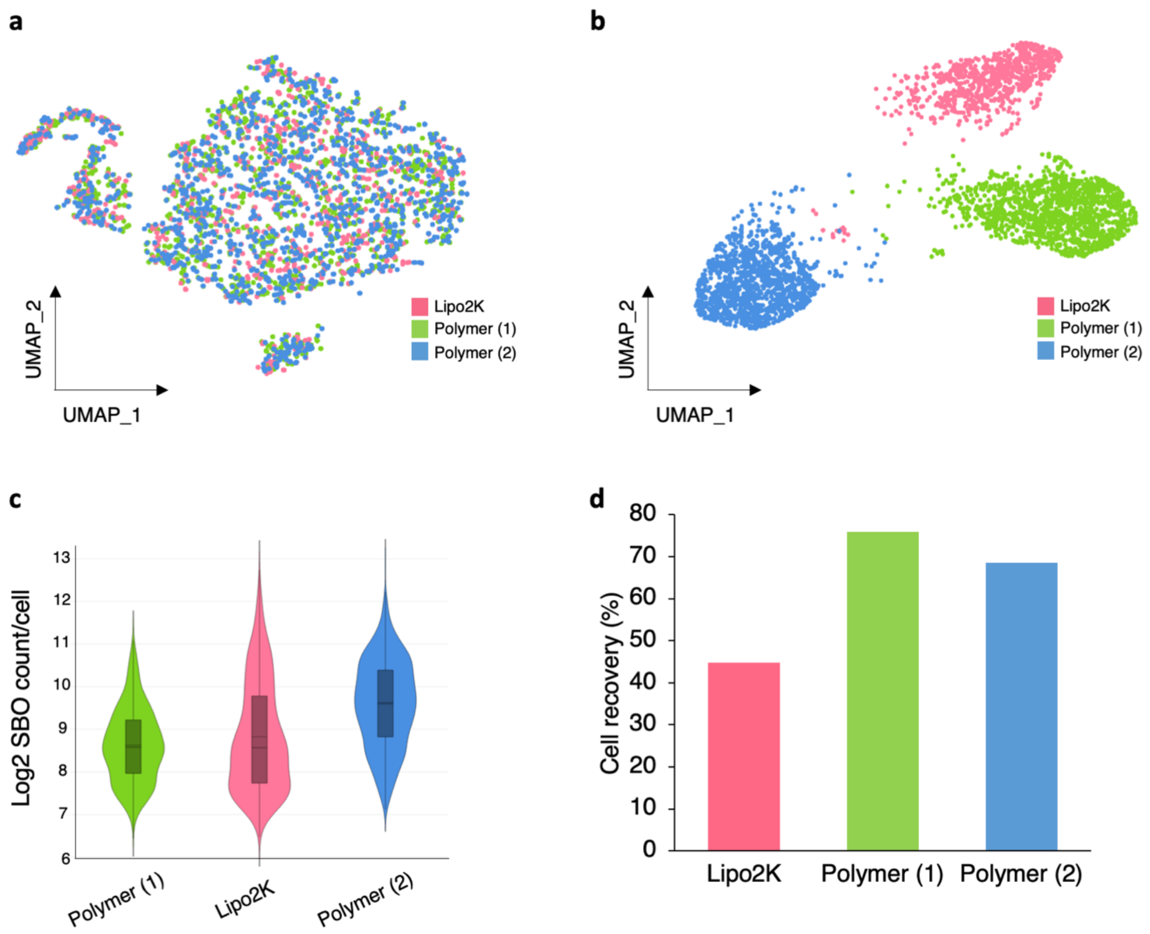


**Supplementary Figure S3. Polymer performs better than Lipofectamine2000 (Lipo2K) at delivering SBOs for multiplex experiments.** Transfection agent indicated by colour; Lipo2K (pink), and polymer (green and blue). Polymer (1) and (2) indicate replicates. (a) UMAP plot of 3-plex HeLa cells. (b) UMAP plot showing the distinct separation of SBOs. (c) $Log_2$ SBO count per cell. (d) Percentage of cells recovered for each sample.

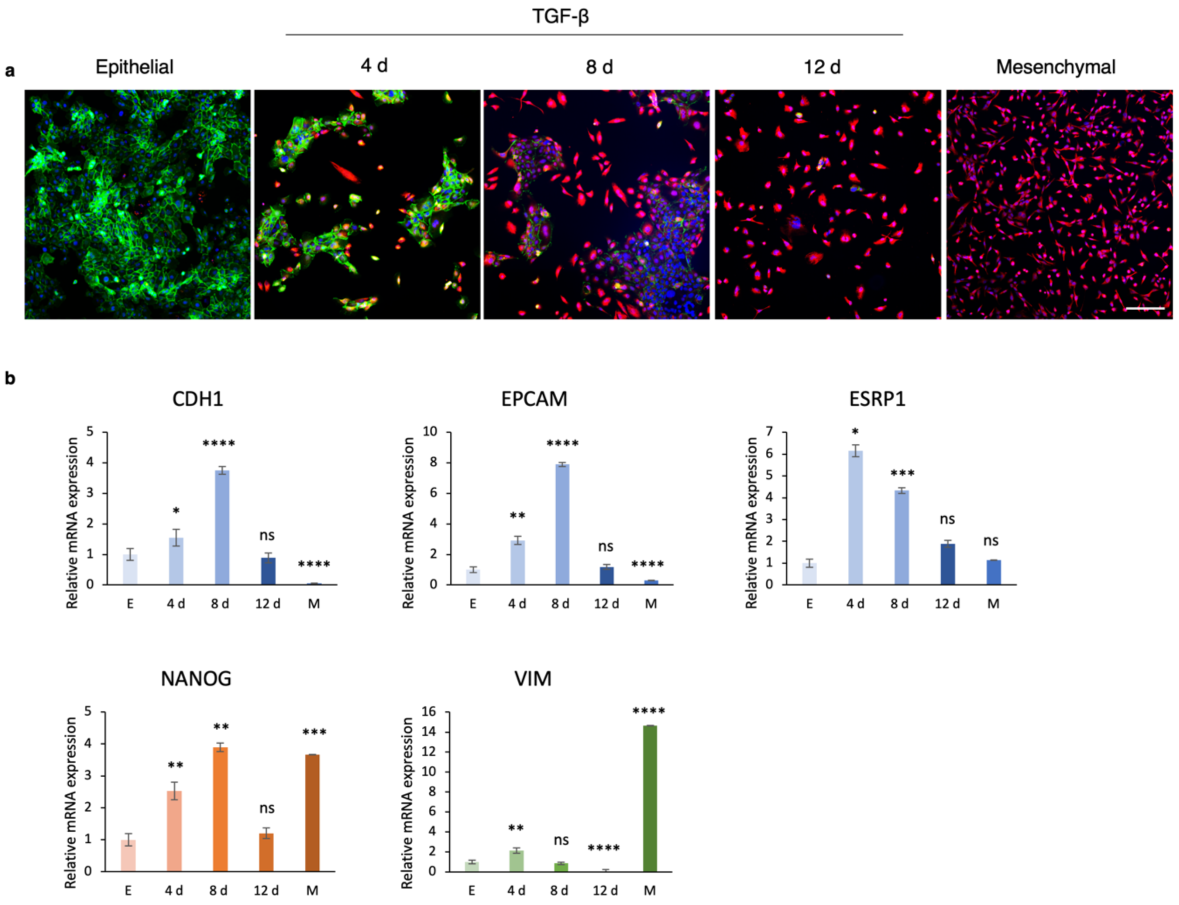


**Supplementary Figure S4. TGF-β stimulated EMT in HMLE cells, resulting in the loss of epithelial markers and gain of mesenchymal markers.** Epithelial HMLE cells (E) were treated with 2 ng/mL TGF-β over 12 days where samples were collected at 4, 8 and 12 days and compared with stable mesenchymal cells mesHMLE (M). (a) Immunofluorescence of epithelial protein E-cadherin (green) and mesenchymal protein vimentin (red) in TGF-β treated HMLE cells (blue nuclei) and mesHMLE cells. (b) Relative gene expression profiles of epithelial genes (CDH1, EPCAM and ESRP1), stemness marker (NANOG) and mesenchymal gene (VIM) in TGF-β treated HMLE cells and mesHMLE cells. Significance calculated using $\Delta\Delta C_t$ values ± S.D. $p$**** < 0.0001, $p$*** < 0.001, $p$** < 0.01, *$p$ < 0.05, ns = non-significant.

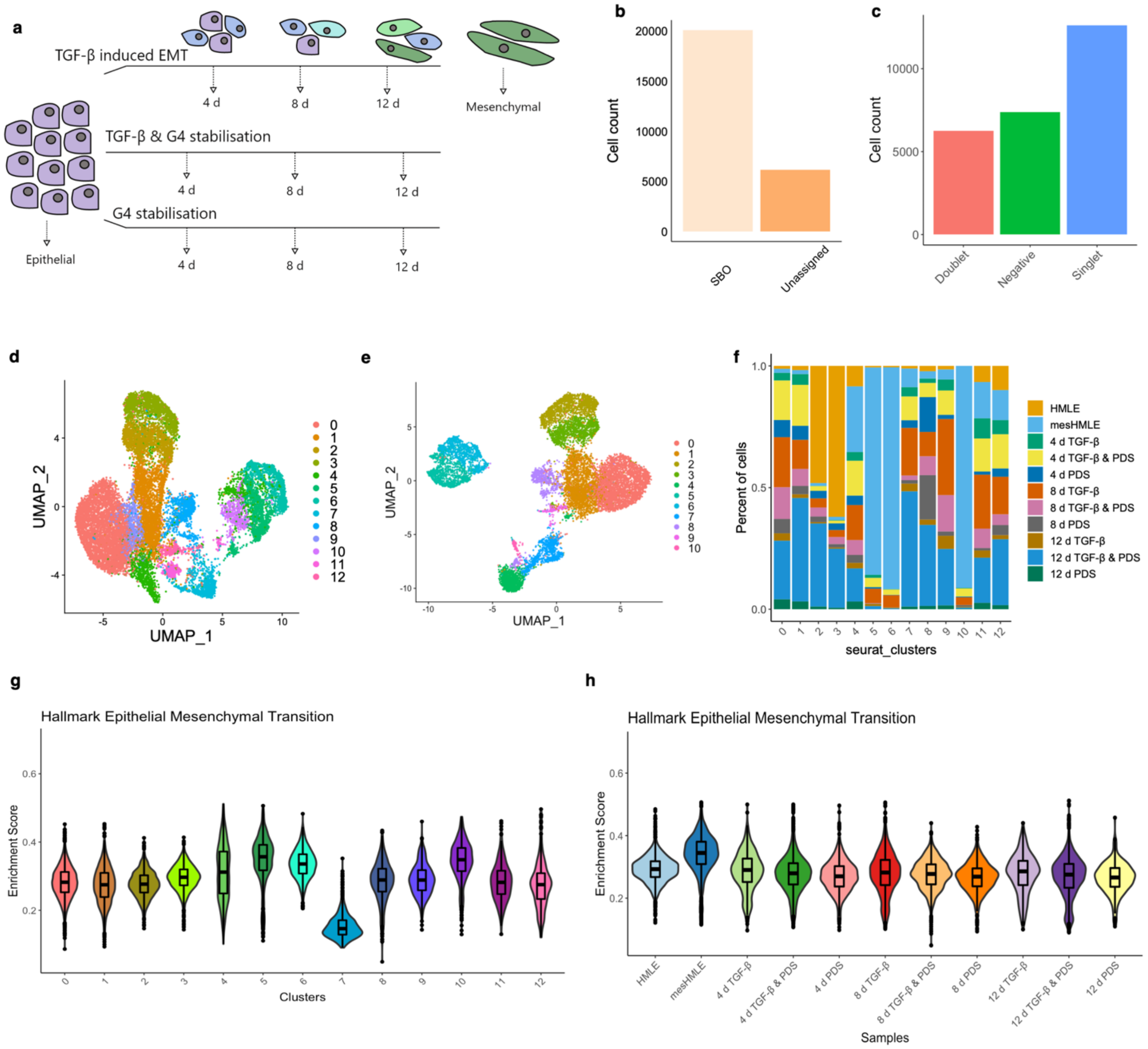


**Supplementary Figure S5. scTECH-seq captures diversity in 12-plex EMT experiment. (a)** Schematic of samples pooled for scTECH-seq to investigate G4 stabilisation on EMT via treatment with Pyridostatin (PDS). **(b)** 76% single cells were assigned a single SBO, with 23% cells unassigned (either doublets or negatives) using the D-score. **(c)** 48% single cells were assigned a single SBO using Seurat HTODemux. **(d)** UMAP clustering of all 11 samples using D-score method of SBO assignment. Clusters are annotated by colour. **(e)** UMAP clustering of all 11 samples using Seurat demultiplexing for SBO assignment. Clusters are annotated by colour. **(f)** Distribution of cells in each cluster. Samples are annotated in colour. **(g - h)** Box and violin plots showing the EMT pathway enrichment for clusters (g) and samples (h).

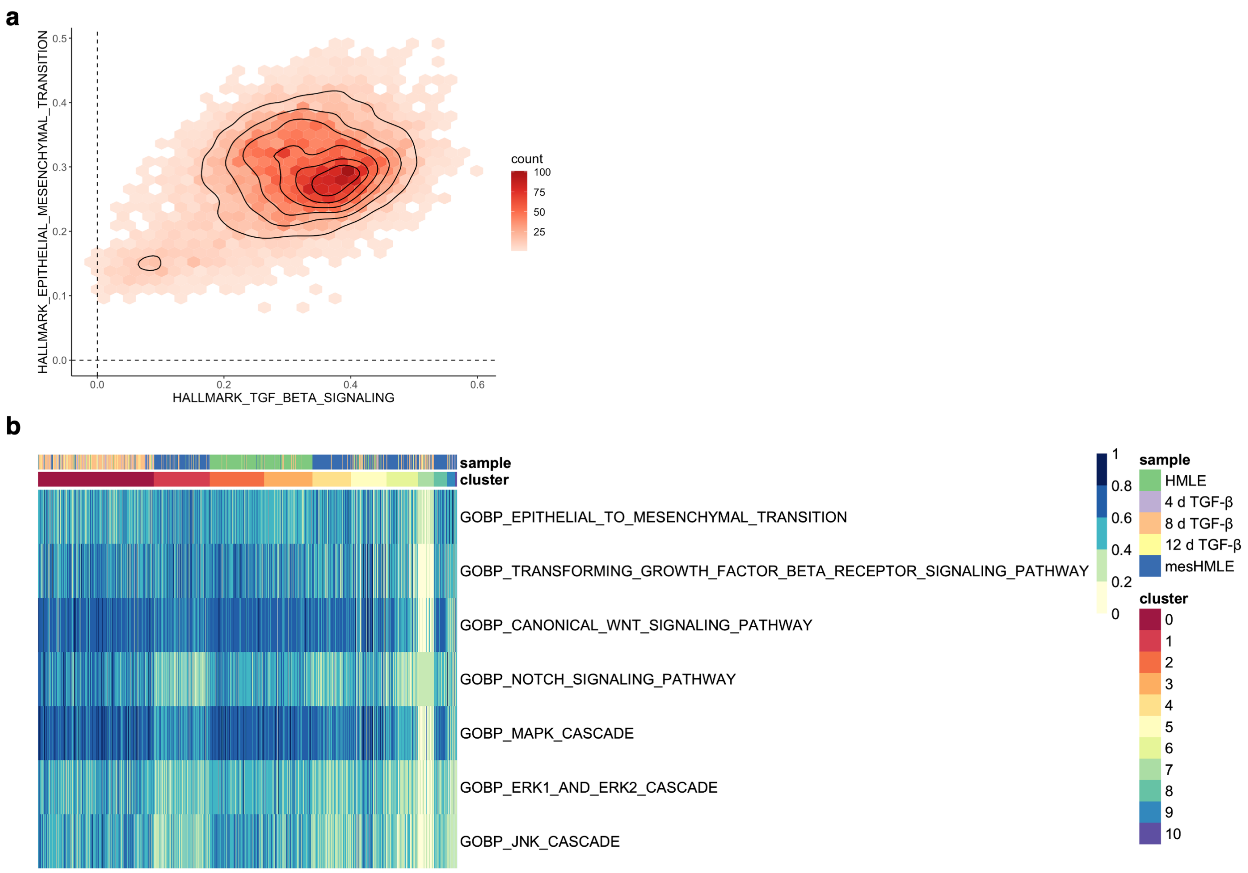


**Supplementary Figure S6. EMT signalling pathways are enriched in TGF-β induced EMT.** Pathway enrichment was determined using ssGSEA R package. **(a)** Cell density of epithelial-to-mesenchymal transition and TGF-β signalling pathway enrichment. Colour scale is indicative of cell count. **(b)** Expression levels of EMT signalling pathways. Each column represents a single cell. Cells are grouped by cluster, with sample assignment shown at the top of the plot.

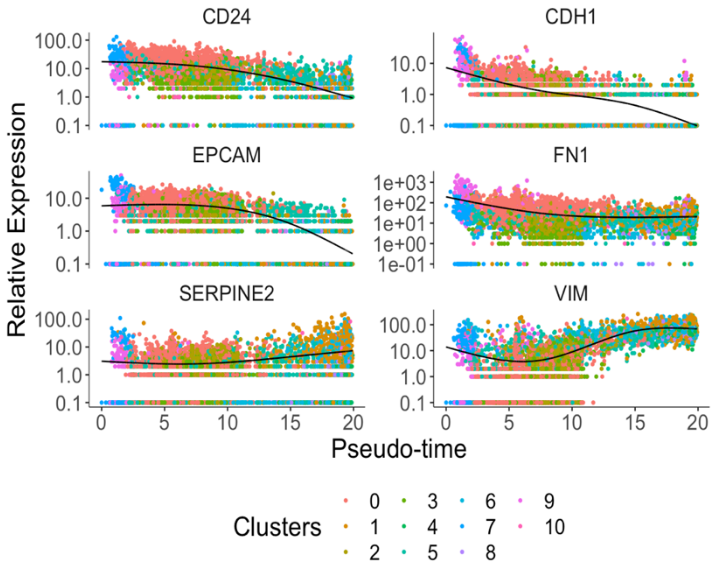


**Supplementary Figure S7. Gene expression alters with EMT pseudotime.** Gene expression alterations in TGF-β EMT. Expression of epithelial genes (*CD24*, *CDH1* and *EPCAM*) decrease with pseudotime, whereas expression of mesenchymal genes (*FN1, SERPINE2* and *VIM*) increase over pseudotime. Cells are annotated by cluster assignment in colour.

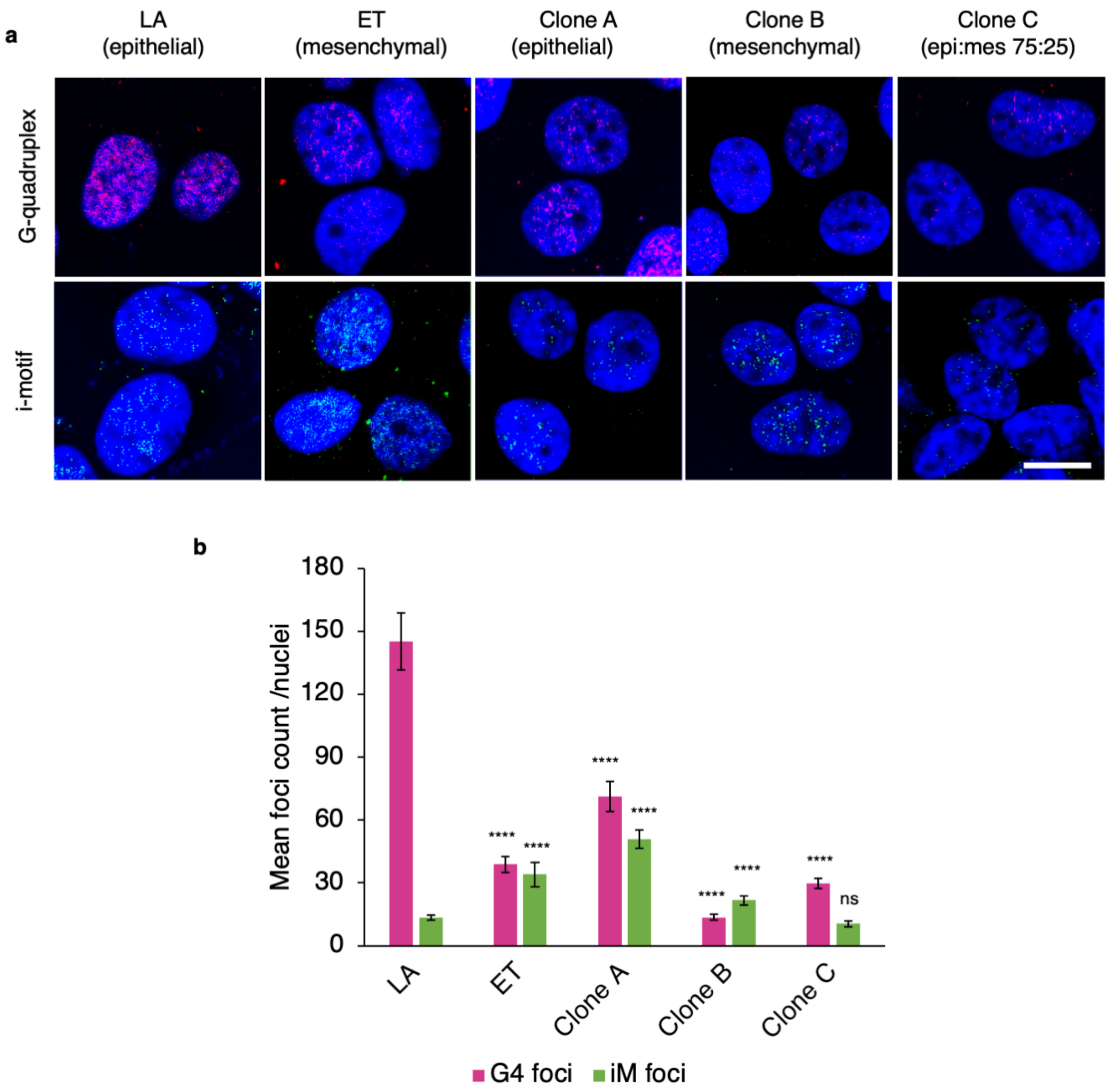


**Supplementary Figure S8. G4 and iM immunofluorescence in PMC42 cell nuclei. (a)** Representative images of G4 (pink) and iM (green) immunofluorescence. Scale bar; 20 µm. **(b)** Quantification of G4 and iM foci per nucleus. $n = 52 - 128$ nuclei were counted per condition. Data displayed as mean ± S.D. Statistical significance shown is relative to PMC42-LA for each structure, ns = not-significant, $p^{***} < 0.001$, $p^{*****} < 0.0001$.

**Supplementary Table S2. Intracellular pH (pHi) of HMLE cells changes with TGF-β stimulation.** Cells were treated with 2 ng/ml TGF-β. Fluorescence was measured at Ex/Em = 490/535 nm. Values shown are the mean from three technical replicates.

| | **HMLE** | **mesHMLE** | **4 d** | **8 d** | **12 d** |
|---|---|---|---|---|---|
| pHi | 7.09 | 7.11 | 8.23 | 8.16 | 7.87 |

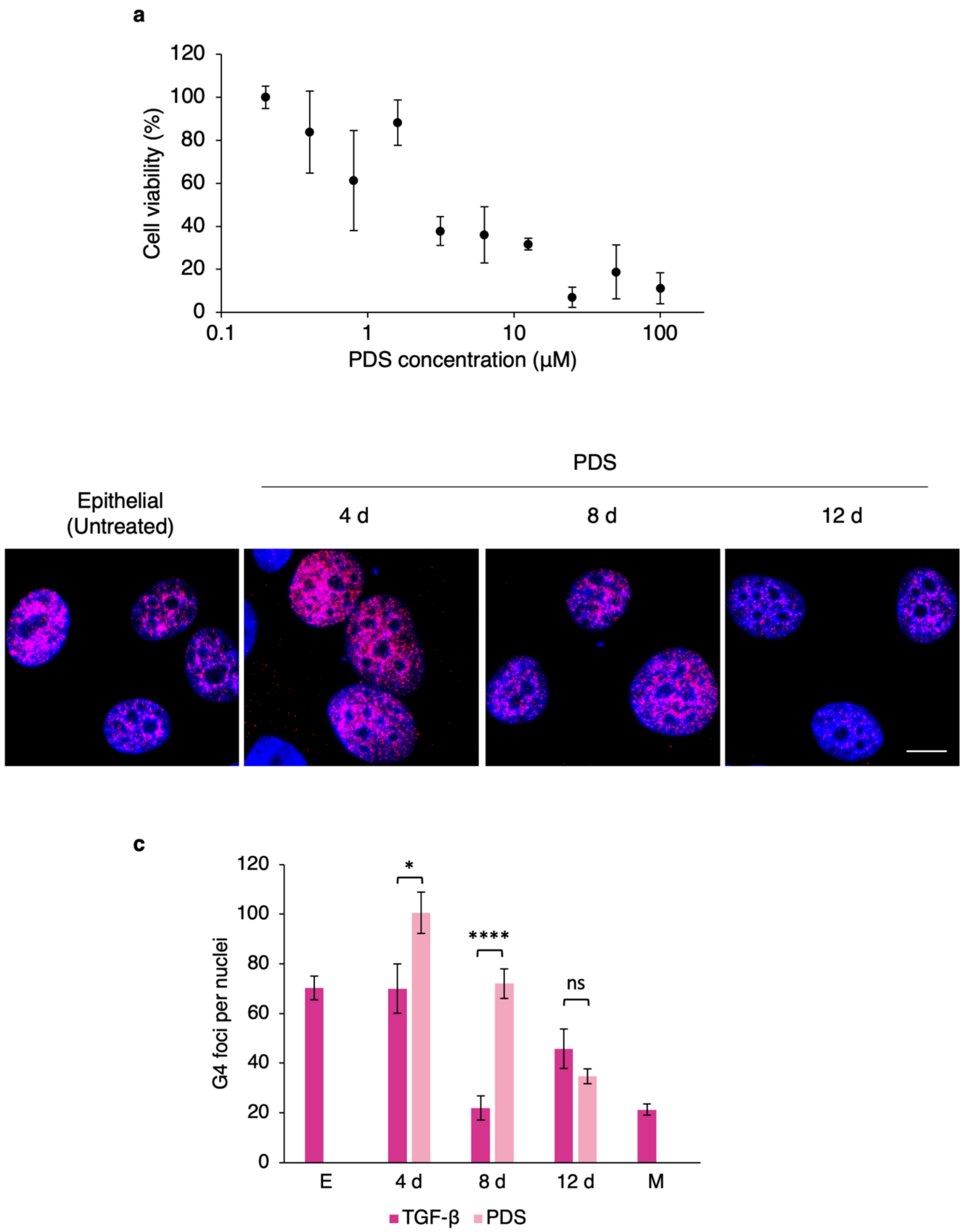


**Supplementary Figure S9. Pyridostatin (PDS) stabilises G4 structures. (a)** Cell viability assay HMLE cells treated with PDS (0.2–100 µM) for 72 h. Data is displayed as mean ± S.D. calculated by three technical replicates. **(b)** Representative images of G4 immunofluorescence (pink foci) in HMLE cells (epithelial). HMLE cells were treated with 0.4 µM PDS for 4, 8, and 12 days. Scale bar, 20 µm. **(c)** Comparison of G4 quantification between TGF-β treated cells (data adapted from Fig. 5b) and PDS treated cells. Data displayed as mean ± S.D. Statistical significance shown; $^{***}p < 0.001$, $^{*}p < 0.05$, ns: $p > 0.05$.

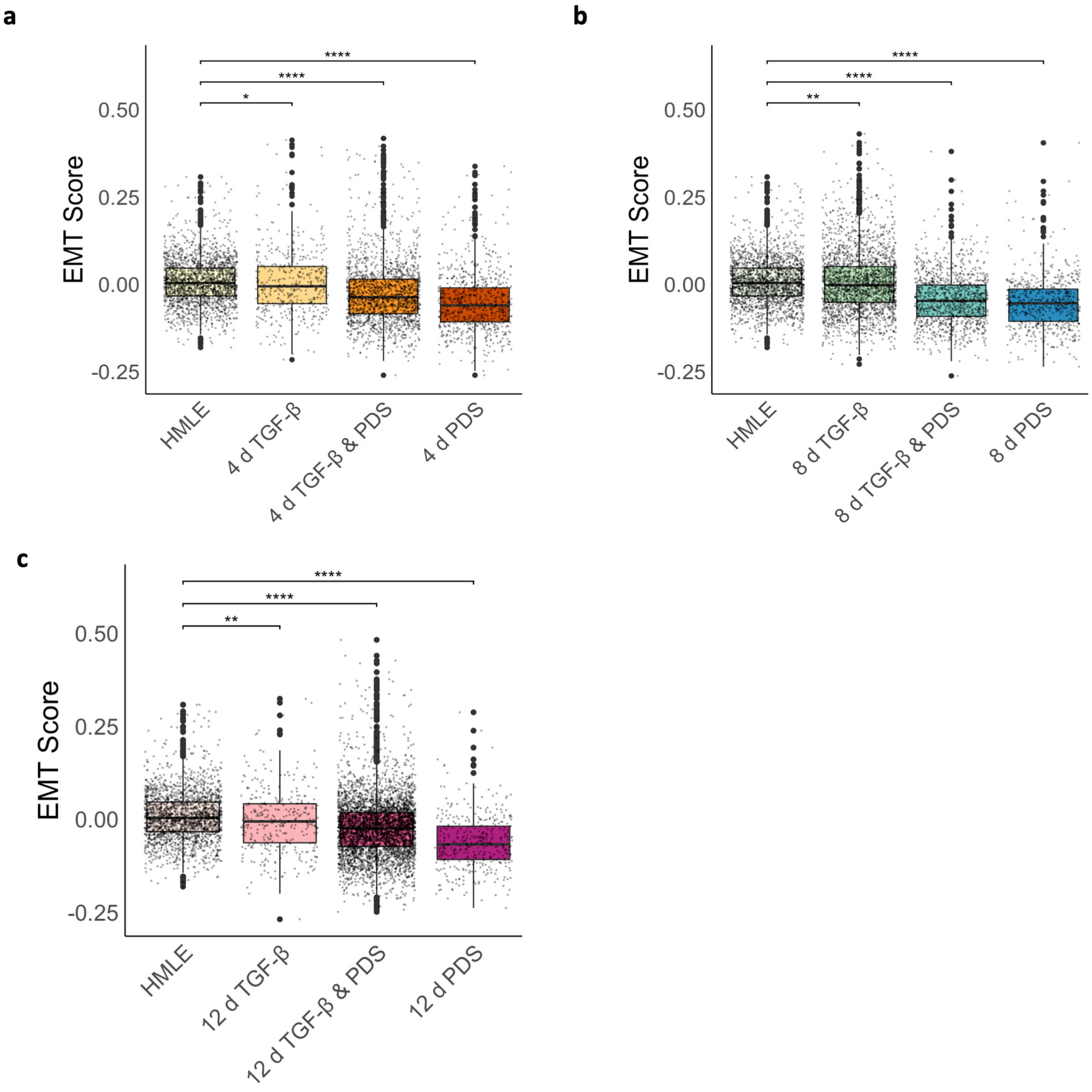


**Supplementary Figure S10. PDS lowers the EMT score of samples.** PDS treatment results in epithelial-like EMT score. HMLE cells treated for four **(a)**, eight **(b)**, and twelve **(c)** days. Boxplots show minimum, first quartile, median, third quartile and maximum EMT score. Individual cell scores are indicated by grey dots and cells that are outliers are displayed as dots. Significance is shown relative to HMLE cells. *$p < 0.05$, **$p < 0.01$, ***$p < 0.001$, ****$p < 0.0001$.

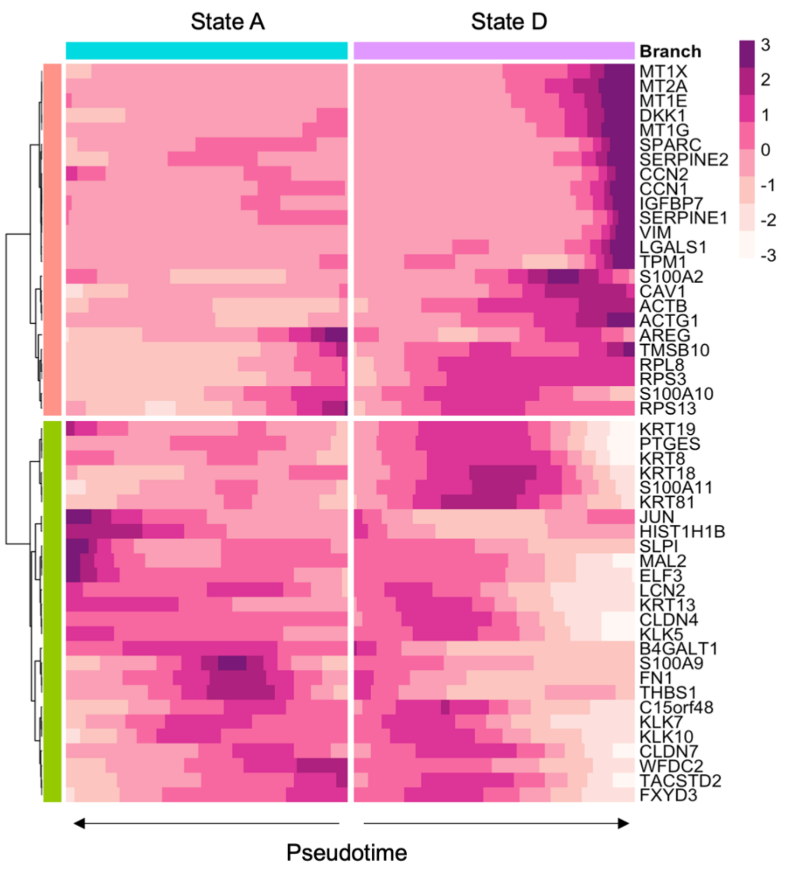


**Supplementary Figure S11. TGF-β and PDS treatment alters gene expression profiles.** Expression heatmap of genes altered with pseudotime in cell state A (pseudotime = 0) and state D (pseudotime = 16).

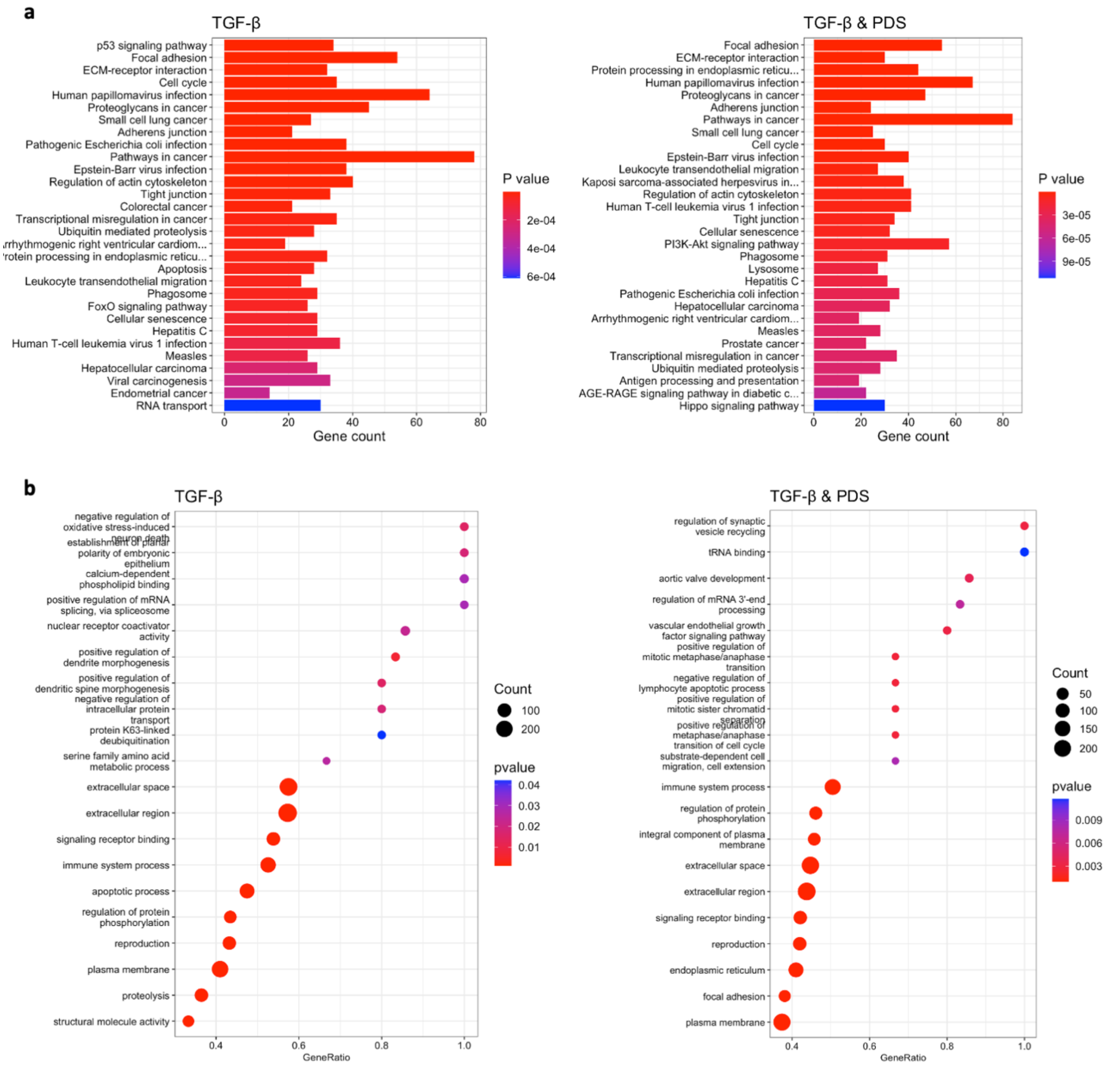


**Supplementary Figure S12. G4 stabilisation alters gene ontology in EMT. (a)** KEGG gene set enrichment in TGF-β EMT with the absence (left) and presence (right) of G4 stabilisation. The top 2,000 differentially expressed genes ($p < 0.05$, $\log_2FC < 0.25$) for each condition were used in enrichment analysis. **(b)** Gene ontology enrichment (biological processes, molecular function and cellular component) in the absence (left) and presence (right) of G4 stabilisation in TGF-β EMT were calculated using ClusterProfiler.

### Supplementary Methods

Polymer characterisation

The dendritic polymer was synthesised outlined by Kretzmann et al.[12] using a 25% GMA backbone and generation 5.0 poly(amido amine) dendrons. Polymer-SBO binding was assessed by a gel retardation assay. Polymer solutions were mixed with 1 µg SBO at different amine-to-phosphate (N/P) ratios and incubated at 25 °C for 30 min. Samples were electrophoresed on a 1% w/v agarose gel in sodium borate buffer at 120 V for 80 min. For dynamic light scattering (DLS) and zeta potential measurements, various N/P polymer ratios were mixed with SBO (1 µg) and incubated for 30 min. Solutions were diluted to 400 µL and measurements were taken in triplicate. Reported values are the mean ± SD for average peak size and zeta potential.

PMC42 cell culture

PMC42 clonal cells were cultured in DMEM (Gibco, cat #11966025) supplemented with 10% v/v FBS and 1x penicillin-streptomycin (Gibco, cat #15140122). Cells were maintained at 37 °C in a humidified incubator with 5% $CO_2$ and passaged at 80% confluency using 1× TrypLE.

Intracellular pH

Abcam Intracellular pH assay (ab228552) was used to measure intracellular pH of cells treated with 2 ng/mL TGF-β as per manufacturer's instructions for standard assay. Fluorescence was measured at ex/em 490/535 nm.
